# Stomach-brain coupling indexes a dimensional signature of mental health

**DOI:** 10.1101/2024.06.05.597517

**Authors:** Leah Banellis, Ignacio Rebollo, Niia Nikolova, Micah Allen

## Abstract

Visceral rhythms orchestrate the physiological states underlying human emotion. Chronic aberrations in these brain-body interactions are implicated in a broad spectrum of mental health disorders. However, the relationship of gastric-brain coupling to affective symptoms remains poorly understood. We investigated the relationship between this novel interoceptive axis and mental health in 243 participants, using a cross validated machine learning approach. We find that increased fronto-parietal brain coupling to the gastric rhythm indexes a dimensional signature of worse mental health, spanning anxiety, depression, stress, and well-being. Control analyses confirm the specificity of these interactions to the gastric-brain axis. Our study proposes coupling between the stomach and brain as a factor in mental health and offers potential new targets for interventions remediating aberrant brain-body coupling.

## Main

Far from being a mere brain in a vat, the nervous system is embedded within an intricate web of visceral rhythms. While philosophy has long championed the embodiment of mind and life^1^, it is only more recently that the importance of the visceral body in contextualising brain function has gained widespread recognition^2,3^. In particular, interoceptive processes linking brain and body are thought to be critically important in mood and emotion^2,4^, and their larger role in affective symptoms has become a topic of intense interest in mental health research^5^.

Research investigating these links have so far focused almost exclusively on the cardiac^6,7^, lower gastrointestinal^8,9^, and respiratory axes^10–12^. Recent landmark findings demonstrate that, for example, anxiety responses in threatening situations relies on ascending cardiac information^6^, and that respiratory rhythms modify neural patterns during emotional processing^13^. Simultaneously, the burgeoning study of gut-brain interaction has produced a plethora of new findings linking the function and biology of the lower gastrointestinal tract or gut to physical and mental health^14–16^. While these and many other findings herald a visceral turn in our understanding of the biology of the mind and its disorder, one particular domain of brain-body coupling remains notably understudied: the upper gastrointestinal tract comprising interconnections between the brain and stomach.

Gastric-brain interactions have recently emerged as a novel frontier in interoception research^17–22^. Hormones secreted by the stomach directly regulate hypothalamic mechanisms that govern satiety and hunger^23^. Additionally, the stomach generates its own independent myoelectrical rhythm, in which the interstitial cells of Cajal pace muscular contractions approximately once every 20 seconds^24^. Previously relegated to merely driving mechanical food digestion, recent discoveries indicate that the gastric rhythm is closely linked to ongoing brain activity through reciprocal vagal innervation^25–27^. This link can be directly modulated through techniques such as non-invasive vagal nerve stimulation^28^, bilateral vagotomy in rodent models^29^, as well as through emerging pharmacological methods^22,30,31^, offering a promising means by which to intervene upon the stomach-brain axis.

Despite the close linkage of emotion and brain-body interaction, the extent to which alterations in the gastric axis relate to mental health remains unclear. This gap is curious in part because folk psychology has long centred the stomach as a locus of stress and anxiety: difficult decisions are described as evoking “gut feelings” which in extreme cases, can make one “sick to their stomach.” Conversely, intense moments of love or joy are described as giving “butterflies in the stomach.” Aligning with these descriptions, recent empirical findings indicate that individuals often report subjective disgust, fear, and anxiety as being localised in the stomach^32,33^, and that pharmacological modulation of the gastric rhythm alters emotional processing^22^. On this basis, we hypothesised that inter-individual patterns in gastric-brain coupling might expose unique patterns of affective mental health, in particular those relating to core mood disorders such as anxiety and depression. To test this hypothesis, we conducted a large scale neuroimaging study of simultaneous electrogastrographic (EGG) and functional MRI (fMRI) brain imaging in 243 participants. To assess mental health across a broad spectrum, we employed a multidimensional approach to quantify highly individualistic profiles spanning a range of affective, cognitive, social, and somatic health dimensions. This approach builds on research identifying mental illness as a continuum of overlapping symptoms across disorders^34–39^, rather than relying on discrete diagnostic categories with high comorbidity, high heterogeneity and poor reliability^40,41^. Utilising a multivariate, cross-validated machine learning technique, we estimated highly robust, sensitive, and specific signatures that interrelate these profiles to individual patterns of stomach-brain coupling. Our findings demonstrate that the stomach-brain axis exposes a positive-to-negative mode of mental health^37,42^, revealing a previously unknown embodied target for future clinical intervention research.

## Results

### Mental health functional correlates

To characterise variation in the gastric-brain axis, we utilised simultaneous electrogastrographic (EGG) and resting state fMRI recordings. Following an extensive data quality control procedure (*Methods)*, we estimated stomach-brain coupling for each participant using the phase locking value (PLV) of the EGG and resting state fMRI time series across 209 parcellated whole-brain regions^43^ (see Figure 1 for example).

**Figure 1:**
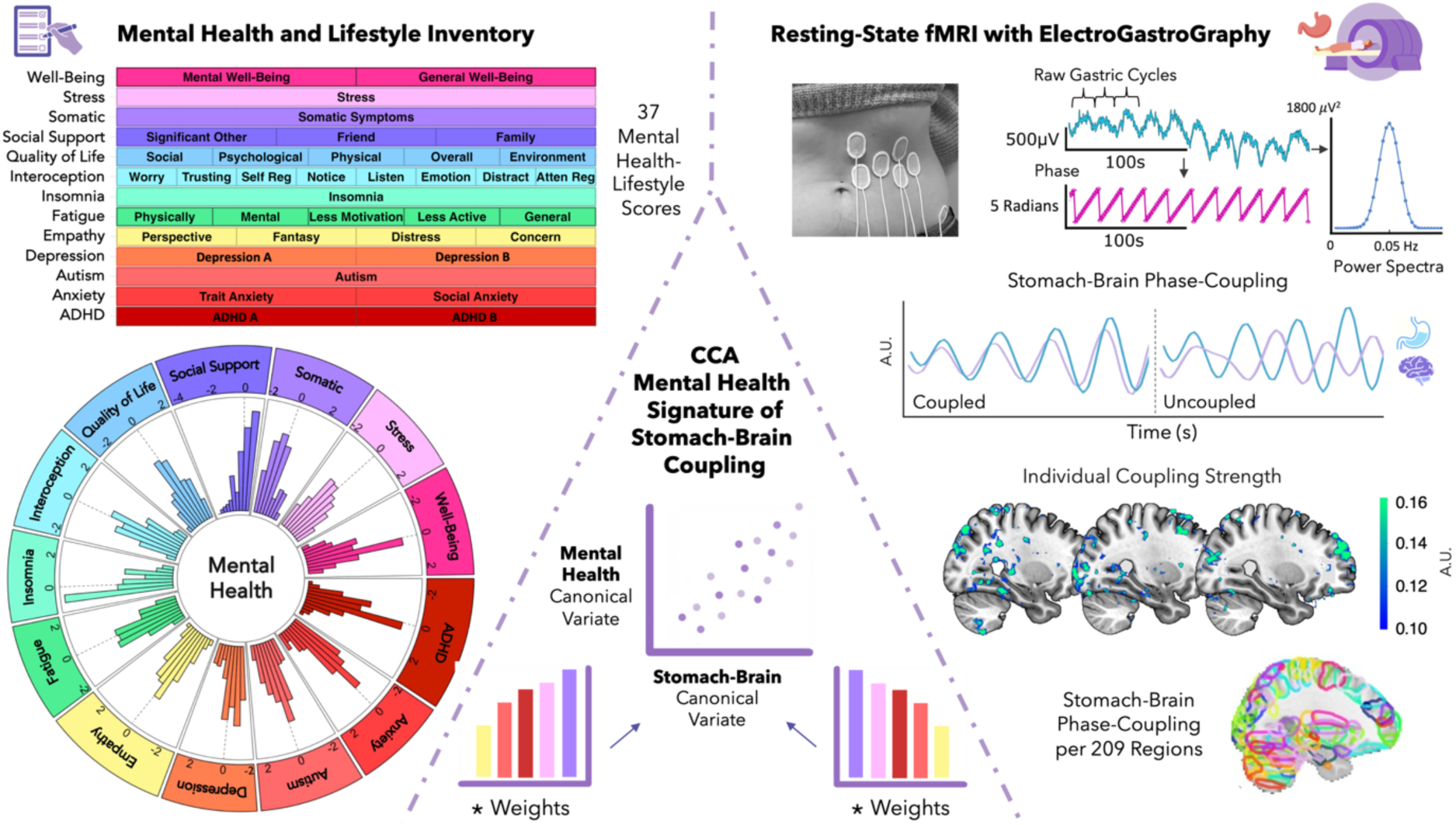
Canonical Correlation Analysis of stomach-brain coupling and mental health. Figure 1 synthesises the process and outcomes of correlating stomach-brain phase coupling with mental health, as quantified by 37 variables from 16 validated surveys. The top left quadrant presents these variables organised into their respective mental health categories (categorised for visualisation only, the CCA incorporated 37 individual scores), and their distribution is visualised as histograms on the bottom left, reflecting the range of participant mental health profiles. Electrogastrography (EGG) data depicted on the top right demonstrates the extraction of gastric cycle frequency from raw EGG signals, power spectra, and their phase information, essential for identifying stomach-brain coupling. The middle right figure illustrates coupled versus uncoupled states in stomach-brain interaction, with the individual variability in coupling strength highlighted across three brain images from individual participants (plotted on a standard mni152 brain template using MRIcroGL: visualised MNI coordinates plotted: 28, −19, 26, thresholded at 0.1, and small clusters <1000mm^3^ removed). For the CCA, stomach-brain phase-coupling is parcellated over 209 brain regions identified using the DiFuMo atlas, shown on the bottom right. The CCA model, depicted centrally, outputs a stomach-brain signature correlating with mental health individual profiles. This pattern is represented by canonical variates, which are weighted combinations of the multidimensional mental health and stomach-brain coupling data (illustrated as the central scatter plot). These weights, depicted as bar graphs, capture the most significant relationships between gastric-brain coupling and mental health profiles.

To characterise individual mental health profiles, we conducted a comprehensive assessment across 37 self-reported scores encompassing autism, ADHD, empathy, insomnia, interoception, depression, fatigue, social support, somatic symptoms, stress, social anxiety, trait anxiety, well-being, and quality of life (see Supplementary Table 1 for a full list of instruments and subscales). This approach successfully captured robust inter-individual variability spanning a variety of mental health dimensions (see Figure 1).

Individual variance spanned from completely healthy to those experiencing significant distress, such that 30% of the sample exhibited mild depression, 19% exhibited clinically significant levels of ADHD, 19% medium or more severe somatic symptoms, 18% trait anxiety, 9% moderate depression, 7% autism spectrum, and 5% insomnia (see Supplementary Table 1 for all percentage cut-offs).

Finally, to determine latent patterns interlinking these mental health profiles to stomach-brain coupling, we conducted a cross-validated Canonical Correlation Analysis (CCA). This method determines maximally correlated patterns between two multidimensional variables (in this case, mental health and stomach-brain coupling data). CCA does this via linear transformation of the inputted data using weights, which produces the resulting CCA variates (i.e., weighted sums) (see Figure 1).

We observed a significant latent dimension in which stomach-brain coupling was associated with a positive-to-negative mode of mental health (canonical variate in-sample *r* (118) = 0.886, out-sample *r* (77) = 0.323, *p* = 0.001). This was reflected behaviourally as high negative loadings for trait anxiety (STAI trait subscale: −0.827), depression (PHQ9: −0.800, and MDI: −0.782), stress (PSS: −0.773), and fatigue (MFI general fatigue subscale: −0.734), as well as high positive loadings for well-being, and quality of life (highest loadings: WEMWBS: 0.856, WHOQOL psychological subscale: 0.847, WHO5: 0.776). The top stomach-brain coupling canonical loadings were found in the left superior angular gyrus (−0.317), right intermediate primus of Jensen (right supramarginal gyrus posterior division using the Harvard-Oxford Cortical Structural Atlas) (−0.284), left inferior precentral sulcus (−0.270), left posterior superior frontal gyrus (−0.269), and left posterior intraparietal sulcus (−0.245). These loadings constitute a pattern of stomach-brain coupling in which healthier mental scores (i.e., improved well-being and quality of life) are associated with reduced gastric coupling to fronto-parietal brain activity (see Figure 2 and Supplementary Table 2), or conversely, in which more negative mental health scores (i.e., increased anxiety, depression, and fatigue) are associated with increased coupling. We did not observe significant gender or age differences with this mental health associated stomach-brain coupling result (see Supplementary Table 3).

**Figure 2:**
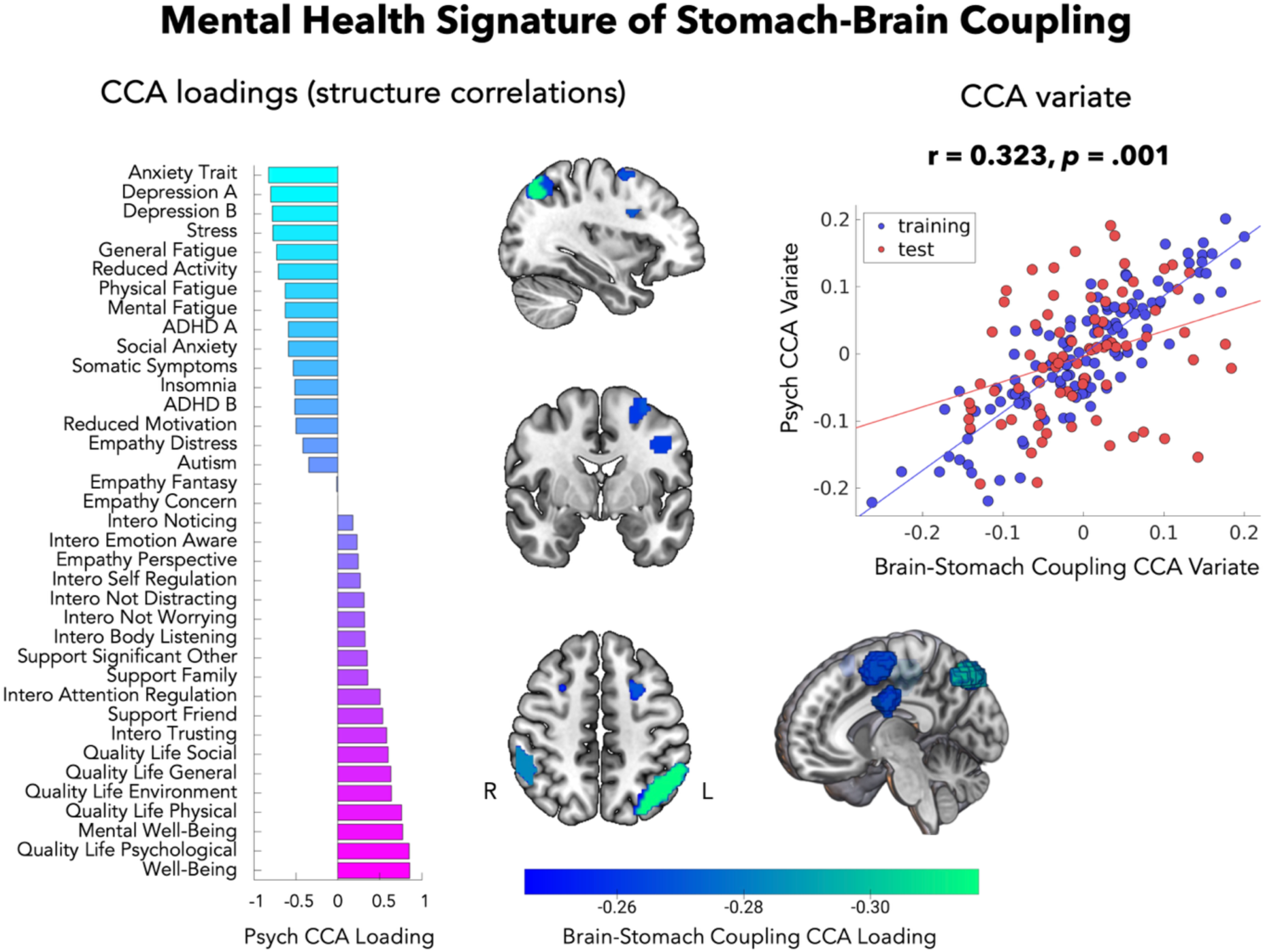
Mental health functional correlate of stomach-brain coupling. Canonical Correlation Analysis results depicting the correlation between stomach-brain coupling and mental health dimensions. This indicated diminished fronto-parietal stomach-brain coupling with healthier mental health scores (i.e., lower anxiety, depression, and stress, and higher quality of life and well-being). Left panel depicts the CCA loadings (structure correlations: Pearson’s correlations between raw mental health and stomach-brain coupling variables and their respective canonical variate). Importantly, this represents the pattern of mental health data that is maximally correlated with the stomach-brain coupling canonical variate. High negative loadings are shown for anxiety, depression, stress, fatigue, ADHD, somatic symptoms, and insomnia, while high positive loadings are shown for well-being and quality of life. The middle panel shows the top 5 DiFuMo parcellated regions with the absolute highest stomach-brain coupling loadings (all negative), coloured according to their respective CCA loading: left superior angular gyrus, right posterior supramarginal gyrus, left inferior precentral sulcus, left posterior superior frontal gyrus and left posterior intraparietal sulcus (plotted on a standard mni152 brain template using MRIcroGL: MNI coordinates: −34, −3, 48). Right depicts the cross-validated CCA result denoting the maximally correlated psychological variate and brain-stomach coupling variate (in-sample *r* (118) = 0.886, out-sample *r* (77) = 0.323, *p* = 0.001).

To further summarise our findings, we averaged parcel-level stomach-brain canonical loadings across the Yeo 7-networks^44^, and also averaged psychological canonical loadings across mental health categories (to condense the multivariate variables, and to be consistent with the mental health categorisation in Figure 1). This revealed the highest absolute network level stomach brain loadings in the dorsal attention (−0.132), frontoparietal control (−0.100), and ventral attention salience network (−0.061). This pattern of stomach-brain coupling was maximally correlated with the corresponding averaged mental health structure, in particular with strong negative loadings for depression (−0.791), stress (−0.773), anxiety (−0.709), and fatigue (−0.642), and strong positive loadings for well-being (0.815) and quality of life (0.696) (see Figure 3, and Supplementary Figure 1 for an averaged summary of the raw weights).

**Figure 3:**
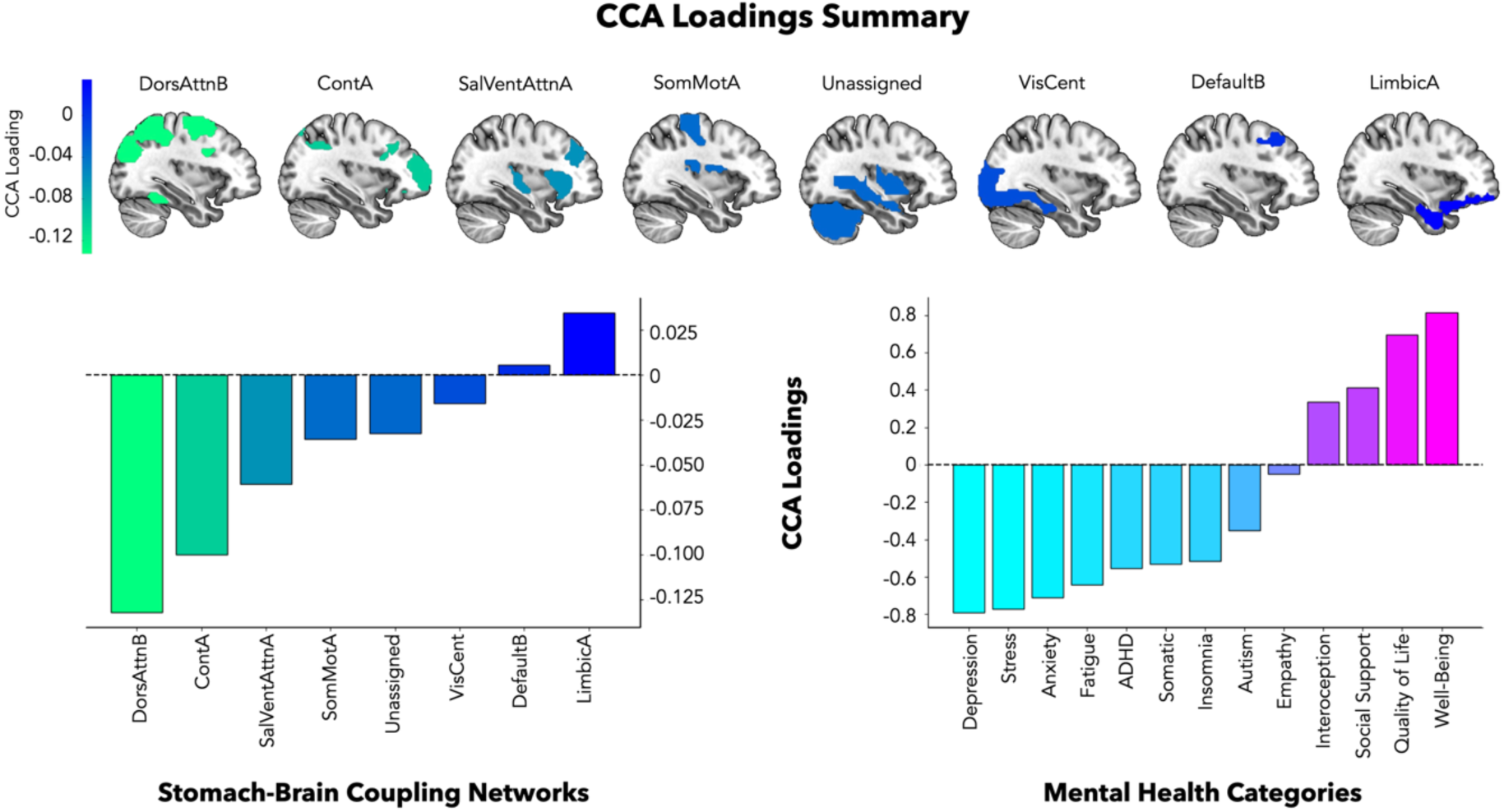
CCA loadings averaged summary. Canonical loadings (structure correlations: Pearson’s correlations between raw inputted variables and respective canonical variate) from the mental health associated stomach-brain coupling CCA, summarised via averaging. Note that there are prominent negative average stomach-brain loadings in the ʻdorsal attention B’ network and the ʻcontrol A’ network, associated with reduced average depression, stress, anxiety, fatigue, and increased average well-being and quality of life (i.e. better mental health). The opposite pattern is also true: increased average stomach-brain loadings in attention and control networks is associated with worse mental health (increased depression, stress, anxiety, fatigue, and reduced well-being, and quality of life). Left shows the stomach-brain loadings averaged according to yeo-7 networks. Above demonstrates these network-averaged stomach-brain loadings projected onto a mask of the DiFuMo regions for each yeo-7 network (from left to right: DorsAttnB = Dorsal Attention B, ContA = Control A, SalVentAttnA = Salience Ventral Attention A, SomMotA = Somatomotor A, Unassigned = no network found, VisCent = Visual A, DefaultB = Default Mode B, LimbicA = Limbic A)^44^, plotted on a standard mni152 brain template using MRIcroGL: MNI coordinates: −34, −3, 48. Right illustrates the psychological loadings averaged across mental health categories defined for visualisation in Figure 1.

### CCA control analyses

To evaluate the specificity and robustness of these results, we conducted control CCAs predicting mental health scores from either 1) functional connectivity, 2) BOLD signal variability, 3) cardiac or 4) respiratory brain maps instead of from stomach-brain coupling. In all cases no significant canonical variate was found (p > 0.05, Bonferonni threshold = 0.01), indicating that the dimensional index of mental health reported here is specific to stomach-brain coupling and unlikely to be explained by residual BOLD connectivity, signal variability or cardiac/respiratory-brain influences.

We conducted a further control analysis to determine if a simpler univariate model would yield comparable results as to our multivariate approach. A principal components analysis (PCA) of the mental health scores yielded a highly similar latent structure as that derived from the CCA (r (35) = −0.999, *p* < .001) (see Supplementary Figure 2). We then correlated this mental health PCA component with stomach-brain coupling values from each DiFuMo parcellated region separately. This revealed significant correlations of stomach-brain coupling with the mental health PCA component in 9 DiFuMo parcellated regions, all of which were in the top loading regions of the CCA, with the exception of the ʻlateral fissure anterior’ (see Supplementary Table 4). Furthermore, across all brain regions, loadings from the stomach-brain coupling CCA were highly correlated with the univariate correlation coefficients linking mental health to stomach-brain coupling (r (207) = −0.870, *p* < .001) (see Supplementary Figure 2). These results complement our cross-validation procedure by demonstrating that the multivariate CCA model detected effects could be reproduced with a simpler, albeit less sensitive, model, with the CCA explaining more variance (in-sample R^2^ = 0.785, out-sample R^2^ = 0.104) than the significant univariate correlations (mean R^2^ = 0.033).

We also controlled for whether the mental health associated stomach-brain coupling result was driven by gastrointestinal symptoms by removing the somatic symptoms survey (PHQ15) from the CCA, and instead including the PHQ15 survey as a nuisance regressor. The CCA result persisted without the somatic symptoms survey with highly replicable loadings (see Supplementary Figure 3). Furthermore, when completing the CCA with the somatic symptom survey items only, or only the gastrointestinal symptom items only, the results were not significant (all somatic symptom survey items smallest p = 0.428, gastrointestinal symptom survey items only smallest p = .267).

### EGG control analyses

To control for possible low-level physiological confounds, we further estimated the association between the psychological canonical variate loadings and summary electrogastrographic (EGG) metrics, using non-parametric Spearman’s rank order correlation coefficients (see Supplementary Table 5 for EGG metric descriptive statistics). We did not observe a significant correlation of normogastric EGG activity measured via the proportion of normogastric power, maximum power, or peak frequency with the observed mental health canonical variate (smallest *FDR-corrected p* = 0.267). Therefore, the link between stomach-brain coupling is specific to the strength of brain-body coupling with mental health, rather than being explained by baseline differences in peripheral gastric physiology.

## Discussion

Our study reveals a distinctive stomach-brain signature of mental health, established through cross-validated multivariate regression techniques and control analyses. This signature encapsulates a positive-to-negative latent dimension of mental health, with notable negative loadings on anxiety, depression, stress, fatigue, as well as enhanced well-being and quality of life. Significantly, all 20 of the highest loadings for stomach-brain coupling were negative, indicating a direct correlation between diminished gastric-brain coupling and improved mental health (refer to Supplementary Table 6). Our control analyses confirm that this extensive psychological signature spanning affective, cognitive, social, and somatic health dimensions is uniquely attributable to stomach-brain coupling, distinct from factors such as residual brain connectivity or variability, cardiac-brain or respiratory-brain influences, gastric activity variations, bodily mass, age, or gender differences. However, future research should explore this using a more diverse sample with a balanced representation of age groups and genders.

Our main result reveals a pattern of worse mental health with increased gastric coupling in brain regions which are known transdiagnostic hotspots, such as the posterior superior frontal gyrus and the posterior intraparietal sulcus^45,46^. Notably, the left superior angular gyrus, our model’s most prominently featured region, is crucial for its integrative role in various cognitive functions^47^. This region is also associated with a range of psychiatric disorders, including schizophrenia^48^, somatization disorder^49^, and major depressive disorder^50,51^. Thus, gastric rhythms may co-vary with brain activity in neural hubs that are highly sensitive to disruptions in mental health. Note, recent advancements have discovered numerous transdiagnostic biotypes in affective disorders like depression and anxiety with varying resting and emotion-evoked connectivity and brain activation profiles^52^. Our findings indicate that there may be brain-body biotypes, however further causal research is necessary. In our study, the brain areas in which psychological health dimensions were most significantly associated with the gastric axis comprised attentional and control network hubs^44^. This indicates that top-down attentional and inhibitory control mechanisms may be particularly important for the relationship of visceral-brain rhythms with mental health. Beyond these control oriented networks, we also observed a negative association with the ventral salience network^53^, weak negative loadings in the somatomotor network^54^, and weak positive loadings in the limbic system, emphasising the multidimensional nature of the signature.

This research significantly advances our understanding of the mental health implications of stomach-brain coupling^18,19,55,56^. While previous smaller scale studies have linked stomach-brain coupling with bodily shame and weight preoccupation^57^, our multivariate CCA approach leverages our extensive sample size to encompass a continuum of transdiagnostic mental health scores. Indeed our multidimensional mental health variate is comparable to previously observed positive-to-negative axes of wellbeing across cognitive, affective, and lifestyle dimensions^37,42^, as well as observed ʻgeneral mental health factors’ which encompass symptoms across numerous psychiatric disorders^35,38,58^. Importantly, although our analyses do not focus on clinical diagnostic categories, our multivariate, psychological dimension-based approach is advantageous in directly assessing highly individualistic mental health profiles across a broad multidimensional spectrum. This aligns with recent paradigm shifting calls for a dimensional schema in mental health with biological plausibility^34,40,41^, as dichotomous psychiatric diagnoses are plagued with short-comings, including arbitrary thresholds for binarisation, poor reliability, high rates of diagnosis comorbidity, shared symptomatology across disorders, and symptom heterogeneity within disorders^40,41^. Moreover, the cross-validated method we apply here is specifically optimised to estimate the continuous statistical prediction of these dimensions, while also robustly protecting against overfitting^59^. Future work could build upon these results to predict multidimensional psychiatric symptoms based on stomach-brain coupling in controlled clinical samples or longitudinal studies.

Interestingly, we identified trait anxiety as a prominent mental health feature associated with stomach-brain coupling, but a previous study found no such relationship with state anxiety^19^. A key difference between that study and ours, was the use of a region-of-interest based approach in a smaller sample size. Our whole-brain, multivariate method was optimised to detect such effects, and likely yielded a substantial improvement in statistical power by estimating latent psychological dimensions directly. It may also be that there are distinct stomach-brain relationships with trait and state aspects of anxiety. Notably, the anxiogenic relationship of stomach-brain coupling is also consistent with a previous report linking state anxiety with intestinal-brain coupling in the insula^60^ and rodent research of increased anxiety behaviours when activating gut-innervating vagal afferents^14^, as well as research with generalised anxiety disorder patients demonstrating an increased bodily reactivity and intensity of interoceptive sensations in response to adrenergic stimulation^7^. Furthermore, a recent pilot study revealed that stress increased gastric phase-amplitude coupling with EEG activity, in contrast to a relaxing biofeedback task^61^. Future research could similarly causally manipulating anxiety or stress to help determine its influence on stomach-brain coupling in fMRI. Complementary approaches could also directly modulate stomach-brain coupling using various emerging interventions as a potential means to remediate anxiety or stress symptoms in patients^17,22,31^.

One potential limitation of our study concerns our EGG data exclusion rate, which was somewhat higher than the 20% rate reported in previous EGG literature^28,62^. This increase was driven by an exhaustive quantitative and qualitative quality control protocol, which may have resulted in higher numbers of excluded participants than in previous, smaller scale studies. To assuage these concerns, we conducted additional control analyses demonstrating the validity of these procedures (see Supplementary Figure 4). Furthermore, we ensured that the excluded participants did not differ in terms of mental health characteristics, gastrointestinal symptoms, or under/over eating behaviour, ruling out the possibility that our quality control may have created a sampling bias which could impact our results (see Electrogastrography Methods). While our preprocessing pipeline aligns with prior literature^18,19,55,56^, emerging ICA-based methods offer promising alternatives for noise reduction, particularly in datasets with high-density montages. Future work may benefit from applying such approaches to reconstruct the EGG signal from components with a high signal-to-noise ratio in the normogastric range^63^.

There is a growing body of evidence associating a dysfunctional gastrointestinal system to various mental health conditions^15,64,65^, as well as on the frequent coexistence of gastrointestinal dysfunction with affective disorders^15^. By elucidating the multimodal interactions between the stomach and the brain in mental health, our findings provide a starting point for future research on novel diagnostic and therapeutic strategies targeting disordered brain-stomach interactions. This includes not only innovations like non-invasive vagus nerve stimulation, which recent studies have found to modulate stomach-brain coupling^28^, but also the exploration of new mechanical^17,66^ and pharmacological^22,31^ interventions to remedy aberrant stomach-brain interactions. Similarly, future research could leverage newly emerging technologies such as ingestible recording devices to further elucidate the physiological mechanisms linking mental health to the stomach-brain axis^67^.

In summary, our study represents the largest and most comprehensive neuroimaging sample focusing on brain-body interaction to date. Our results signify a link between increased stomach-brain coupling and poorer mental health across anxiety, depression, stress, and well-being dimensions. This finding contributes significantly to multidisciplinary research on brain-body interaction and opens new avenues for therapeutic, diagnostic, and classification strategies to improve psychological well-being and mental health.

## Methods

### Participants

We recruited participants as part of the Visceral Mind Project, a large brain imaging project at the Centre of Functionally Integrative Neuroscience, Aarhus University. We recorded electrogastrography (EGG) in 380 participants (230 females, 149 males, 1 other gender, median age = 24, age range = 18-56). As our aim was to apply machine learning to individual differences in mental health, we adopted a participant recruitment strategy that sought to maximise between individual variance from fully healthy to those with scores crossing clinical thresholds. Accordingly, we did not explicitly exclude participants for psychiatric diagnosis, and recruited participants from a wide range of possible online communities and backgrounds (see Supplementary Table 1). These participants did not report any major physical illnesses, or medication beyond over-the-counter antihistamines or contraceptives, furthermore they reported abstinence from alcohol/drugs 48 hours before participation. We acquired participants in two data collection cohorts by advertising on nation-wide participant pools, social media, newspapers, and posted fliers. As additional criteria, participants had normal or corrected-to-normal vision and were fluent in Danish or English. Furthermore, we only included participants compatible with MRI scanning (not pregnant/breastfeeding, no metal implants, claustrophobia etc).

Participants took part in multiple sessions including fMRI scans, behavioural tasks, physiological recordings, and mental health/lifestyle inventories. In this study, we focus on resting state fMRI data, electrogastrography (EGG) recordings, and a mental health/lifestyle assessment battery to evaluate individual differences in gastric-brain coupling and their link to mental health. We compensated individuals for participating. The local Region Midtjylland Ethics Committee granted approval for the study and all participants provided informed consent. The study was conducted in accordance with the Declaration of Helsinki. After the removal of poor-quality fMRI and EGG data (see quality control below), we estimated stomach-brain-coupling in 243 participants. Including the mental health scores, a total of 199 full-dataset participants were included in the mental health functional correlate analysis (138 females, 61 males, median age = 23, age range = 18-47) (see Supplementary Figure 5 and Supplementary Figure 6).

### Anatomical and resting state fMRI acquisition

We acquired anatomical MRI and resting state fMRI data using a 3T MRI scanner (Siemens Prisma) with a 32-channel head coil. We positioned small cushions around the head to minimise head movement. The participants wore earplugs and were instructed not to move. The resting state scan included 600 volumes acquired over 14 minutes using a T2*-weighted echo-planar imaging (EPI) multiband accelerated sequence (TR = 1400ms, TE = 29.6ms, voxel size = 1.79 × 1.79 × 1.80 mm). An acceleration factor of 4 was used in the slice direction along with GRAPPA in-plane acceleration factor = 2. A set of high-resolution whole brain T1-weighted anatomical images (0.9 mm^3 isotropic) were acquired using an MP-RAGE sequence (repetition time = 2.2s, echo time = 2.51ms, matrix size = 256 × 256 × 192 voxels, flip angle = 8°, AP acquisition direction).

### Physiological recording acquisition

We simultaneously recorded physiological measurements (photoplethysmography, respiratory breathing belt, and EGG) during resting-state fMRI. For the EGG recordings, we cleaned the abdomen and applied abrasive gel to remove dead skin and improve the signal-to-noise ratio. Three electrogastrography recording montages were implemented using a Brain Vision MRI-compatible ExG system and amplifier (see Supplementary Figure 7 for each recording montage consisting of 1, 3 or 6 bipolar channels). All physiological montages were acquired with a sampling rate of 1000 Hz, a low-pass filter of 1000 Hz (with a 450 Hz anti-aliasing filter), and no high-pass filter (DC recordings). EGG was recorded at a 0.5 µV/bit resolution, and +/− 16.384 mV range, while photoplethysmography and respiratory recordings were acquired at 152.6 µV/bit resolution, and +/− 5000 mV range.

### MRI and fMRI preprocessing

We implemented the minimal preprocessing pipeline in fmriprep. MRI and fMRI results included in this manuscript come from preprocessing performed using fMRIPrep 22.1.1, which is based on Nipype 1.8.5 (see Supplementary Material for anatomical and functional MRI preprocessing details with fMRIPrep). Additional fMRI preprocessing steps following fMRIPrep included spatial smoothing with a 3mm FWHM kernel, and regressing out six motion parameters, six aCompCor parameters, as well as 13 RETROICOR components reflecting cardiac and respiratory physiological noise.

### Electrogastrography peak selection and preprocessing

The EGG data was first demeaned and downsampled from 1000 Hz to 10 Hz for computational efficiency, followed by computing the power spectrum using a Hanning-tapered fast Fourier transform (FFT) incorporating 1000 seconds of zero-padding in 200-second data segments with 75% overlap. For each participant, we selected the bipolar EGG channel that showed the most prominent peak within the normal frequency range of the gastric rhythm in humans (i.e., normogastric range: 0.033-0.066 Hz), which is on average one cycle every 20 seconds (0.05 Hz)^62^. Specifically, two researchers (L.B. and I.R.) independently conducted peak selection by visually inspecting each channel to identify the EGG channel with the highest normogastric power peak, without large artefacts and with power above 5 µv^2^. Peak quality was rated as ‘excellent’ for gaussian-like peaks (n=184) and ‘good’ for shoulder-like peaks (n=81); those not meeting these standards were deemed ‘poor quality’ (n=115) and excluded. This visual inspection approach is consistent with previous research to account for noise in the normogastric window, or cases of multiple peaks^18,19,28^. Note, selected electrode choice did not cause significant differences in the mental health CCA variate (F range(2 to 5, 86 to 87) = 0.867 to 1.972, p range = .146 to .507, n^2^ range = 0.044 to 0.051) or the stomach-brain coupling CCA variate (F range(2 to 5, 86 to 87) = 1.376 to 2.863, p range = .063 to .242, n^2^ range = 0.063 to 0.078). Similarly, the EGG recording montage did not cause any significant differences in the mental health CCA variate (F(2, 198) = 1.717, p = .182, n^2^ = 0.017) or the stomach-brain coupling CCA variate (F(2, 198) = 1.220, p = .298, n^2^ = 0.012).

As an additional check, we computed signal quality metrics using a comparison template-based procedure of 10 ideal participants with very clear and prominent gastric peaks. ʻPoor quality’ participants had significantly lower signal quality as measured by cosine similarity (excellent/good quality: Median = 0.963, Range = 0.667, poor quality: Median = 0.595, Range = 0.585; U = 63780, *p* < .001, r_rb_ = 3.186) and Pearson’s correlation (excellent/good quality: Median = 0.950, Range = 1.169, poor quality: Median = 0.054, Range = 1.287; U = 63849, p < 0.001, r_rb_ = 3.190) (see Supplementary Figure 4). As an extra precaution, we confirmed that the mental health scores of the included and excluded EGG participants did not significantly differ when using the first PCA component of the 37 mental health scores (excellent/good quality: Median = 3.766, Range = 150.429, poor quality: Median = 5.745, Range = 106.762; U = 46989, *p* = 0.964, r_rb_ = 2.084). Furthermore, included/excluded participants did not differ in reported gastrointestinal symptoms (average of PHQ15 items inquiring of “stomach pain”, “constipation, loose bowels, or diarrhea”, and “nausea, gas, or indigestion“: excellent/good quality: Median = 1.333, Range = 2, poor quality: Median = 1.333, Range = 2; U = 14104, p = 0.913, rrb = −0.0744). In addition, participants did not differ in reported under/overeating behaviour (PHQ9 item “poor appetite or overeating“: excellent/good quality: Median = 1, Range = 3, poor quality: Median = 1, Range = 3; U = 13250, p = 0.361, rrb = −0.130). The selected EGG channel was then bandpass filtered, centred at the individual peak frequency (filter width of ±0.015 Hz, filter order of 5 or 1470 samples), in forward and backward direction to avoid time shifts. After phase correction, the data was resampled to the fMRI rate (0.7143 Hz) and processed through a Hilbert transform to calculate the average phase per volume.

### Gastric-brain coupling estimation

We followed procedures validated in previous EGG studies to estimate gastric-brain coupling^18,19^. The preprocessing of BOLD time series for all brain voxels involved bandpass filtering, using parameters identical to those applied during the EGG analysis. The initial and final 21 volumes (equivalent to 29.4 seconds) were excluded from both the fMRI and EGG time series. This adjustment resulted in a total signal duration of 781.2 seconds for further analysis. The instantaneous phases of both signals were obtained through the application of the Hilbert transform. Subsequently, the phase-locking value (PLV) was calculated as the absolute value of the average phase angle differences between the EGG and each voxel over time (see Equation 1)^43^. The PLV is quantified by values ranging from 0 (representing a total absence of phase synchrony) to 1 (corresponding to absolute phase synchrony).

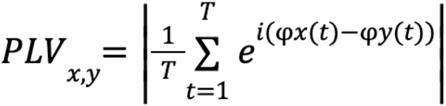

***Equation 1***: where T is the number of time samples, and x and y are brain and gastric time series.

In order to account for any biases in PLV that arise from differences in signal amplitude, we created surrogate PLV values by disrupting the phase relationship between EGG and BOLD time series. We achieved this by shifting the EGG by at least ±60 s with respect to the BOLD time series, with concatenation at the edges. Given the 558 samples in the BOLD time series, this procedure generated 472 surrogate PLV datasets. We then took the median value of these surrogate PLV distributions as chance level coupling, and defined coupling strength as the difference between empirical and chance level coupling. Therefore, a higher value represents stronger stomach-brain coupling strength.

### Mental health assessment battery

Participants completed a battery of mental health and lifestyle assessments. This encompassed 16 separate survey instruments spanning autism, ADHD, empathy, insomnia, interoception, depression, fatigue, social support, somatic symptoms, stress, social anxiety, trait anxiety, well-being, and quality of life. All scales utilised validated Danish translations, except in cases where participants spoke English as their first language, in which case validated English versions were used. This allowed us to explore a broad range of mental health and lifestyle factors across 37 subscale scores (see Supplementary Table 1 for details of surveys, abbreviations, and subscale scores).

### Canonical correlation analysis

We used the CCA-PLS toolbox to fit multivariate, cross-validated Canonical Correlation Analysis (CCA) models relating stomach-brain coupling (coupling strength of the BOLD and EGG time series) to mental health scores. Specifically, CCA aims to find linear combinations of each multidimensional variable (i.e., canonical variates: which are weighted sums of stomach-brain coupling (V = X * B) and mental health (U = Y * A)) that are maximally correlated with each other, but uncorrelated with all other combinations (X and Y represent the inputted stomach-brain coupling and mental health data, while A and B represent the canonical weights)^59,68^. The toolbox incorporates various CCA/PLS models, including the cross-validated and optimised PCA-CCA techniques applied here^59^. This method importantly guards against overfitting via optimised data-reduction methods, assessing statistical inference between independent training and test sets, as well as by implementing permutation testing based on the out-of-sample correlation (see below for further details).

We first reduced the dimensions of stomach-brain coupling per fMRI voxel by parcellating with the 256-region Dictionary of Functional Modes (DiFuMo) atlas, excluding regions of cerebrospinal fluid, ventricles, or white matter, yielding 209 relevant regions. Because CCA is very sensitive to outliers^69–72^, it is important to screen for outliers in the stomach-brain coupling and mental health data, leading to the exclusion of 12 and 25 participants respectively (see Supplementary Figure 5 for a complete flow chart of exclusions). This avoids false dependencies in the training set and distortions to the canonical projection weights^69–72^. Our final Canonical Correlation Analysis (CCA) sample comprised 199 participants for whom we had complete stomach-brain coupling and mental health matrices. These were standardised to have zero mean and unit variance, and nuisance regression was applied to control the estimated canonical variates for the influence of gender, age, body mass index, and data collection cohort.

Subsequently, we applied the cross-validated CCA approach within the predictive framework (machine learning) provided by the CCA-PLS toolbox^59^. This predictive approach involved randomly splitting the data into a training/optimisation set (60% of the overall data) and a test/holdout set (40% of the overall data) 5 times (5-fold cross-validation). These optimisation and holdout sets are known as the ʻouter data splits’, used for statistical inference (determining the number of significant associative CCA modes). The p-values were calculated via permutation testing (1000 permutations), as the fraction of the shuffled permuted out-of-sample correlations exceeding the out-of-sample correlation in the non-permuted holdout set. Because we implemented 5 holdout sets, the p-value for each holdout set was Bonferroni corrected (*a* = 0.05/5 = 0.01). An associative CCA effect is considered significant if the p-value was significant in at least one of the independent test/holdout sets, once trained on the training/optimisation set (out-of-sample correlation). If a significant associative CCA effect was found, the CCA iteratively removed the effect from the data via deflation and repeated this approach to find orthogonal CCA associative effects.

Before statistical inference, to overcome multicollinearity and overfitting issues, the PCA-CCA approach optimises the number of features (PCA components) inputted to each of the outer data splits used for statistical inference. Thus, the PCA-CCA approach further divides the optimisation set into a training set (60% of the optimisation set) and a validation set (40% of the optimisation set) 5 times (5-fold hyperparameter optimisation) for each outer data split. These ʻinner data splits’ were used to select the optimal hyperparameters (number of PCA components the inputted stomach-brain coupling and mental health data dimensions were reduced to) by maximising the average out-of-sample correlation in the validation sets.

To aid interpretation of the networks underlying the estimated brain-stomach coupling signature, we visualised overall network-level contributions by averaging the canonical loadings across the Yeo 7-network parcellation for the stomach-brain coupling axis^44^. Moreover, we averaged across mental health categories for the psychological loading axis for clearer visualisation of the CCA result (see Figure 3).

### Control analyses

To control for underlying influences of neural connectivity or brain activity variability to the mental health stomach-brain coupling result, we conducted two separate control CCA analyses. For the neural connectivity control analysis, we parcellated the fMRI preprocessed data (with an additional high-pass filter to handle low-frequency signal drifts) with the same method used for the stomach-brain coupling data (using the 256-DiFuMo atlas and removing cerebrospinal fluid, ventricles, and white matter regions), and quantified individual functional connectivity data matrices using correlation (Nilearn function: ʻConnectivityMeasure.fit_transform’). Furthermore, for the brain activity variability control analysis, we calculated the standard deviation of BOLD activity for each voxel of the fMRI preprocessed data and parcellated using the same method. Both the resting connectivity and BOLD signal variability control CCA analyses were completed with the same CCA parameters as described in the methods section of the manuscript (including gender, age, body mass index, and data collection cohort as nuisance variables).

To control that the mental health dimension we uncovered is specific to the stomach and not cardiac and respiratory activity, we conducted two additional CCA analyses to control for cardiac-brain and respiratory-brain interactions. For the cardiac domain we used inter-beat-intervals of the heartbeat, computed using identified R-peaks via the ʻppg_peaks’ function from systole which uses a rolling average algorithm, while the respiration analysis focused on inhalation breath durations (inter-breath-intervals), computed using identified inhalation peaks via the ʻfind_peaks’ function from scipy.signal with a distance of 1 samples and a peak prominence of 0.6. Both identified cardiac R-peaks and respiratory inhalation peaks were visually inspected and manually corrected if necessary. To estimate instantaneous HRV regressors, we interpolated the cardiac inter-beat-intervals at the fMRI scanner frequency (TR=1.4 seconds, spline interpolation method) and band-pass filtered them at the frequencies corresponding to low (0.05-0.15 Hz) and high (0.15-0.357 Hz - upper limit constrained by Nyquist frequency of the scanner) heart rate variability (low center frequency = 0.1 Hz ± 0.05, high center frequency = 0.2535 Hz ± 0.1035, Matlab FIR filter)^18,73^. For the respiratory domain, we interpolated the inter-breath-intervals at fMRI scanner frequency, and bandpass filtered at 0.1-0.357 Hz^74,75^ (center frequency = 0.2285 Hz ± 0.1285 - upper limit constrained by Nyquist frequency of the scanner). We obtained the amplitude envelopes of the instantaneous high and low frequency HRV and respiratory rate variability signal via a Hilbert transformation. These amplitude envelopes were used as regressors of interest (without convolution with HRF)^18,73^ in first level GLM’s, with six motion and six acompcor noise regressors using SPM12 and a high-pass filter with cutoff of 128 seconds. The fMRI had the same preprocessing as the stomach-brain phase coupling analysis. We then used T-contrasts to identify individual maps of brain activity associated with increased low frequency HRV, high frequency HRV, or respiratory rate variability. Each of these individual heart/respiratory-brain maps were parcellated and inputted into a CCA with the 37 mental health scores with the same parameters as the stomach-brain coupling CCA.

As an additional control, we conducted a separate whole brain analysis to determine if we could identify a similar result when using a simpler mass univariate analysis. First, we computed PCA on the mental health scores to get a single component similar to the psychological canonical loadings from the CCA. This independent mental health PCA component was then correlated with the stomach-brain coupling from each of the 209 DiFuMo parcellated regions separately. Finally, these univariate Pearson correlation coefficients were correlated with the stomach-brain coupling loadings from the multivariate CCA to determine similarity of the two analysis strategies.

Moreover, we completed control Spearman correlations of gastric physiology (EGG metrics) with the mental health canonical variate extracted from the stomach-brain coupling CCA analysis. From the computed EGG power spectra (as described in the electrogastrography preprocessing section), we quantified the following normogastric EGG metrics: peak frequency, maximum power, and proportion of power (see Supplementary Table 5). Specifically, within the normogastric frequency range (0.033-0.067 Hz/2-4 cpm/15-30 seconds), we stored the peak frequency and maximum power. Furthermore, we computed the proportion of normogastric power as the sum of the normogastric power divided by the sum of the power in all gastric frequencies (including bradygastric, normogastric, and tachygastric frequencies: 0.02-0.17 Hz/1-10 cpm/6-60 seconds). We input those EGG metrics into a correlation matrix with the mental health canonical variate, correcting for multiple comparisons using the false-discovery rate (FDR) at 5%.

To control for age or gender effects, we conducted the mental health associated stomach-brain coupling CCA with the same parameters as the main analysis, but removed age and gender as nuisance regressors. We then tested for age and gender effects via a pearsons correlation with age and an independent-samples t-test with gender with the subsequent CCA variate for each stomach-brain coupling and mental health.

## Data availability

Deidentified participant data and scripts implemented in this paper are available here: https://github.com/embodied-computation-group/StomachBrain-MentalHealth

## Acknowledgements

This research is financially supported by a Lundbeckfonden Fellowship (R272-2017-4345) and a European Research Council Grant (ERC-2020-StG-948788). The funding source was not involved in the study design, collection, analysis, interpretation, or writing of the manuscript.

## Author contributions

LB and IR analysed the data, interpreted the results, and wrote the manuscript, NN provided conceptual advice and contributed towards preprocessing of neuroimaging data, MA provided supervision, conceptual advice, and wrote the manuscript.

## Materials and correspondence

Leah Banellis.

## References

1. Thompson, E. Mind in Life: Biology, Phenomenology, and the Sciences of Mind. (Harvard University Press, 2010).

2. Engelen, T., Solcà, M. & Tallon-Baudry, C. Interoceptive rhythms in the brain. Nat. Neurosci. 26, 1670–1684 (2023).

3. Allen, M. Unravelling the Neurobiology of Interoceptive Inference. Trends Cogn. Sci. 24, 265–266 (2020).

4. Critchley, H. D. & Garfinkel, S. N. Interoception and emotion. Curr. Opin. Psychol. 17, 7–14 (2017).

5. Nord, C. L. & Garfinkel, S. N. Interoceptive pathways to understand and treat mental health conditions. Trends Cogn. Sci. 26, 499–513 (2022).

6. Hsueh, B. et al. Cardiogenic control of affective behavioural state. Nature 615, 292–299 (2023).

7. Teed, A. R. et al. Association of Generalized Anxiety Disorder With Autonomic Hypersensitivity and Blunted Ventromedial Prefrontal Cortex Activity During Peripheral Adrenergic Stimulation: A Randomized Clinical Trial. JAMA Psychiatry 79, 323–332 (2022).

8. Cryan, J. F. & Dinan, T. G. Mind-altering microorganisms: the impact of the gut microbiota on brain and behaviour. Nat. Rev. Neurosci. 13, 701–712 (2012).

9. Kolobaric, A. et al. Gut microbiome predicts cognitive function and depressive symptoms in late life. Mol. Psychiatry 1–12 (2024) doi:10.1038/s41380-024-02551-3.

10. Allen, M., Varga, S. & Heck, D. H. Respiratory rhythms of the predictive mind. Psychol. Rev. No Pagination Specified-No Pagination Specified (2022) doi:10.1037/rev0000391.

11. Bagur, S. et al. Breathing-driven prefrontal oscillations regulate maintenance of conditioned-fear evoked freezing independently of initiation. Nat. Commun. 12, 2605 (2021).

12. Kluger, D. S. et al. Modulatory dynamics of periodic and aperiodic activity in respiration-brain coupling. Nat. Commun. 14, 4699 (2023).

13. Zelano, C. et al. Nasal Respiration Entrains Human Limbic Oscillations and Modulates Cognitive Function. J. Neurosci. 36, 12448–12467 (2016).

14. Krieger, J.-P. et al. Neural Pathway for Gut Feelings: Vagal Interoceptive Feedback From the Gastrointestinal Tract Is a Critical Modulator of Anxiety-like Behavior. Biol. Psychiatry 92, 709–721 (2022).

15. Margolis, K. G., Cryan, J. F. & Mayer, E. A. The Microbiota-Gut-Brain Axis: From Motility to Mood. Gastroenterology 160, 1486–1501 (2021).

16. Foster, J. A. & McVey Neufeld, K.-A. Gut–brain axis: how the microbiome influences anxiety and depression. Trends Neurosci. 36, 305–312 (2013).

17. Mayeli, A. et al. Parieto-occipital ERP indicators of gut mechanosensation in humans. Nat. Commun. 14, 3398 (2023).

18. Rebollo, I., Devauchelle, A.-D., Béranger, B. & Tallon-Baudry, C. Stomach-brain synchrony reveals a novel, delayed-connectivity resting-state network in humans. eLife 7, e33321 (2018).

19. Rebollo, I. & Tallon-Baudry, C. The sensory and motor components of the cortical hierarchy are coupled to the rhythm of the stomach during rest. J. Neurosci. 42, 2202–2220 (2021).

20. Azzalini, D., Rebollo, I. & Tallon-Baudry, C. Visceral Signals Shape Brain Dynamics and Cognition. Trends Cogn. Sci. (2019) doi:10.1016/j.tics.2019.03.007.

21. Teckentrup, V. & Kroemer, N. B. Mechanisms for survival: vagal control of goal-directed behavior. Trends Cogn. Sci. 0, (2023).

22. Nord, C. L., Dalmaijer, E. S., Armstrong, T., Baker, K. & Dalgleish, T. A Causal Role for Gastric Rhythm in Human Disgust Avoidance. Curr. Biol. 31, 629–634.e3 (2021).

23. Inui, A. Ghrelin: An orexigenic and somatotrophic signal from the stomach. Nat. Rev. Neurosci. 2, 551–560 (2001).

24. Sanders, K. M., Ward, S. M. & Koh, S. D. Interstitial Cells: Regulators of Smooth Muscle Function. Physiol. Rev. 94, 859–907 (2014).

25. Richter, C. G., Babo-Rebelo, M., Schwartz, D. & Tallon-Baudry, C. Phase-amplitude coupling at the organism level: The amplitude of spontaneous alpha rhythm fluctuations varies with the phase of the infra-slow gastric basal rhythm. NeuroImage 146, 951–958 (2017).

26. Cao, J. et al. Gastric stimulation drives fast BOLD responses of neural origin. NeuroImage 197, 200–211 (2019).

27. Alladin, S. N. B., Judson, R., Whittaker, P., Attwood, A. S. & Dalmaijer, E. S. Review of the gastric physiology of disgust: Proto-nausea as an under-explored facet of the gut– brain axis. Brain Neurosci. Adv. 8, 23982128241305890 (2024).

28. Müller, S. J., Teckentrup, V., Rebollo, I., Hallschmid, M. & Kroemer, N. B. Vagus nerve stimulation increases stomach-brain coupling via a vagal afferent pathway. Brain Stimulat. 15, 1279–1289 (2022).

29. Cao, J., Wang, X., Chen, J., Zhang, N. & Liu, Z. The vagus nerve mediates the stomach-brain coherence in rats. NeuroImage 263, 119628 (2022).

30. Marso Steven P., et al. Semaglutide and Cardiovascular Outcomes in Patients with Type 2 Diabetes. N. Engl. J. Med. 375, 1834–1844 (2016).

31. Wilding John P.H., et al. Once-Weekly Semaglutide in Adults with Overweight or Obesity. N. Engl. J. Med. 384, 989–1002 (2021).

32. Nummenmaa, L., Hari, R., Hietanen, J. K. & Glerean, E. Maps of subjective feelings. Proc. Natl. Acad. Sci. 115, 9198–9203 (2018).

33. Nummenmaa, L., Glerean, E., Hari, R. & Hietanen, J. K. Bodily maps of emotions. Proc. Natl. Acad. Sci. 111, 646–651 (2014).

34. Insel, T. R. & Cuthbert, B. N. Brain disorders? Precisely. Science 348, 499–500 (2015).

35. Calkins, M. E. et al. The Philadelphia Neurodevelopmental Cohort: constructing a deep phenotyping collaborative. J. Child Psychol. Psychiatry 56, 1356–1369 (2015).

36. Gillan, C. M., Kosinski, M., Whelan, R., Phelps, E. A. & Daw, N. D. Characterizing a psychiatric symptom dimension related to deficits in goal-directed control. eLife 5, e11305 (2016).

37. Smith, S. M. et al. A positive-negative mode of population covariation links brain connectivity, demographics and behavior. Nat. Neurosci. 18, 1565–1567 (2015).

38. Kaczkurkin, A. N. et al. Common and dissociable regional cerebral blood flow differences associate with dimensions of psychopathology across categorical diagnoses. Mol. Psychiatry 23, 1981–1989 (2018).

39. Benwell, C. S. Y., Mohr, G., Wallberg, J., Kouadio, A. & Ince, R. A. A. Psychiatrically relevant signatures of domain-general decision-making and metacognition in the general population. Npj Ment. Health Res. 1, 1–17 (2022).

40. Insel, T. et al. Research Domain Criteria (RDoC): Toward a New Classification Framework for Research on Mental Disorders. Am. J. Psychiatry 167, 748–751 (2010).

41. Schumann, G. et al. Stratified medicine for mental disorders. Eur. Neuropsychopharmacol. 24, 5–50 (2014).

42. Goyal, N., Moraczewski, D., Bandettini, P. A., Finn, E. S. & Thomas, A. G. The positive– negative mode link between brain connectivity, demographics and behaviour: a pre-registered replication of Smith et al. (2015). R. Soc. Open Sci. 9, 201090 (2022).

43. Lachaux, J.-P., Rodriguez, E., Martinerie, J. & Varela, F. J. Measuring phase synchrony in brain signals. Hum. Brain Mapp. 8, 194–208 (1999).

44. Yeo, B. T. T. et al. The organization of the human cerebral cortex estimated by intrinsic functional connectivity. J. Neurophysiol. 106, 1125–1165 (2011).

45. McTeague, L. M. et al. Identification of Common Neural Circuit Disruptions in Cognitive Control Across Psychiatric Disorders. Am. J. Psychiatry 174, 676–685 (2017).

46. Yang, Y. et al. Common and Specific Functional Activity Features in Schizophrenia, Major Depressive Disorder, and Bipolar Disorder. Front. Psychiatry 10, (2019).

47. Seghier, M. L. The Angular Gyrus. The Neuroscientist 19, 43–61 (2013).

48. Gao, Y. et al. Decreased resting-state neural signal in the left angular gyrus as a potential neuroimaging biomarker of schizophrenia: An amplitude of low-frequency fluctuation and support vector machine analysis. Front. Psychiatry 13, (2022).

49. Su, Q. et al. Decreased interhemispheric functional connectivity in insula and angular gyrus/supramarginal gyrus: Significant findings in first-episode, drug-naive somatization disorder. Psychiatry Res. Neuroimaging 248, 48–54 (2016).

50. Gao, J. et al. Habenula and left angular gyrus circuit contributes to response of electroconvulsive therapy in major depressive disorder. Brain Imaging Behav. 15, 2246–2253 (2021).

51. Mo, Y. et al. Bifrontal electroconvulsive therapy changed regional homogeneity and functional connectivity of left angular gyrus in major depressive disorder. Psychiatry Res. 294, 113461 (2020).

52. Tozzi, L. et al. Personalized brain circuit scores identify clinically distinct biotypes in depression and anxiety. Nat. Med. 30, 2076–2087 (2024).

53. Menon, V. & Uddin, L. Q. Saliency, switching, attention and control: a network model of insula function. Brain Struct. Funct. 214, 655–667 (2010).

54. Gordon, E. M. et al. A somato-cognitive action network alternates with effector regions in motor cortex. Nature 617, 351–359 (2023).

55. Choe, A. S. et al. Phase-locking of resting-state brain networks with the gastric basal electrical rhythm. PLOS ONE 16, e0244756 (2021).

56. Levakov, G., Ganor, S. & Avidan, G. Reliability and validity of brain-gastric phase synchronization. Hum. Brain Mapp. 44, 4956–4966 (2023).

57. Todd, J., Cardellicchio, P., Swami, V., Cardini, F. & Aspell, J. E. Weaker implicit interoception is associated with more negative body image: Evidence from gastric-alpha phase amplitude coupling and the heartbeat evoked potential. Cortex 143, 254–266 (2021).

58. Schürmann-Vengels, J., Troche, S., Victor, P. P., Teismann, T. & Willutzki, U. Multidimensional Assessment of Strengths and Their Association With Mental Health in Psychotherapy Patients at the Beginning of Treatment. Clin. Psychol. Eur. 5, e8041 (2023).

59. Mihalik, A. et al. Canonical Correlation Analysis and Partial Least Squares for Identifying Brain–Behavior Associations: A Tutorial and a Comparative Study. Biol. Psychiatry Cogn. Neurosci. Neuroimaging 7, 1055–1067 (2022).

60. Hashimoto, T. et al. Neural correlates of electrointestinography: Insular activity modulated by signals recorded from the abdominal surface. Neuroscience 289, 1–8 (2015).

61. Jeanne, R. et al. Gut-Brain Coupling and Multilevel Physiological Response to Biofeedback Relaxation After a Stressful Task Under Virtual Reality Immersion: A Pilot Study. Appl. Psychophysiol. Biofeedback 48, 109–125 (2023).

62. Wolpert, N., Rebollo, I. & Tallon-Baudry, C. Electrogastrography for psychophysiological research: Practical considerations, analysis pipeline, and normative data in a large sample. Psychophysiology 57, e13599 (2020).

63. Alladin, S. N. B. et al. Children aged 5–13 years show adult-like disgust avoidance, but not proto-nausea. Brain Neurosci. Adv. 8, 23982128241279616 (2024).

64. Chang, L., Wei, Y. & Hashimoto, K. Brain–gut–microbiota axis in depression: A historical overview and future directions. Brain Res. Bull. 182, 44–56 (2022).

65. Vindegaard, N., Speyer, H., Nordentoft, M., Rasmussen, S. & Benros, M. E. Gut microbial changes of patients with psychotic and affective disorders: A systematic review. Schizophr. Res. 234, 41–50 (2021).

66. Rao, S. S. C., Quigley, E. M. M., Chey, W. D., Sharma, A. & Lembo, A. J. Randomized Placebo-Controlled Phase 3 Trial of Vibrating Capsule for Chronic Constipation. Gastroenterology 164, 1202–1210.e6 (2023).

67. Porciello, G., Monti, A., Panasiti, M. S. & Aglioti, S. M. Ingestible pills reveal gastric correlates of emotions. eLife 13, e85567 (2024).

68. Linke, J. O. et al. Shared and Anxiety-Specific Pediatric Psychopathology Dimensions Manifest Distributed Neural Correlates. Biol. Psychiatry 89, 579–587 (2021).

69. Luo, L., Wang, W., Bao, S., Peng, X. & Peng, Y. Robust and sparse canonical correlation analysis for fault detection and diagnosis using training data with outliers. Expert Syst. Appl. 236, 121434 (2024).

70. Wilms, I. & Croux, C. Robust sparse canonical correlation analysis. BMC Syst. Biol. 10, 72 (2016).

71. Okan Sakar, C. & Kursun, O. A method for combining mutual information and canonical correlation analysis: Predictive Mutual Information and its use in feature selection. Expert Syst. Appl. 39, 3333–3344 (2012).

72. Mandal, A. & Cichocki, A. Non-Linear Canonical Correlation Analysis Using Alpha-Beta Divergence. Entropy 15, 2788–2804 (2013).

73. Critchley, H. D. et al. Human cingulate cortex and autonomic control: converging neuroimaging and clinical evidence. Brain 126, 2139–2152 (2003).

74. Natarajan, A. et al. Measurement of respiratory rate using wearable devices and applications to COVID-19 detection. NPJ Digit. Med. 4, 136 (2021).

75. Iqbal, T., Elahi, A., Ganly, S., Wijns, W. & Shahzad, A. Photoplethysmography-Based Respiratory Rate Estimation Algorithm for Health Monitoring Applications. J. Med. Biol. Eng. 42, 242–252 (2022).

