## Supplementary Material for "Stomach-brain coupling indexes a dimensional signature of mental health"

**Supplementary Table 1: Mental Health *and Lifestyle Inventory, including screening diagnosis cutoff percentages for the CCA sample.***

| Survey | Full Name | Scores | Diagnosis Cutoff (%) |
|---|---|---|---|
| **AQ10** (1,2) | Autism Spectrum Quotient | Autism. | 7% *(binarised aq10 sum-score >= 6)* (1) |
| **ASRS** (3) | Adult ADHD Self-Report Scale | ADHD.A (ASRS Part A), ADHD.B (ASRS Part B). | 19% *(binarised asrs.A sum-score >= 4)* (3) |
| **IRI** (4,5) | Interpersonal Reactivity Index | Empathy: Personal-Distress, Fantasy, Empathic-Concern, Perspective-Taking. | NA |
| **ISI** (6) | Insomnia Severity Index | Insomnia. | 5% clinical moderate *(isi sum-score 15-21)*<br>31% subthreshold *(isi sum-score 8-14)* (6) |
| **MAIA** (7) | Multidimensional Assessment of Interoceptive Awareness | Interoception: Noticing, Emotional-Awareness, Self-Regulation, Not-Distracting, Not-Worrying, Body-Listening, Attention-Regulation, Trusting. | NA |
| **MDI** (8,9) | Major Depression Inventory | Depression.B. | 1% moderate *(mdi sum-score 26-30)*<br>9% mild *(mdi sum-score 21-25)* (8) |
| **MFI** (10) | Multidimensional Fatigue Inventory | Fatigue: General-Fatigue, Physical-Fatigue, Mental-Fatigue, Reduced-Activity, Reduced-Motivation. | NA |
| **MPSSS** (11) | Multidimensional Perceived Social Support Scale | Social Support: Significant-Other, Family, Friends. | NA |

| | | | |
|---|---|---|---|
| **PHQ9**<br>(12,13) | Patient Health Questionnaire: depression module | Depression.A. | 9% moderate *(phq9 sum-score 10-14)*<br>30% mild *(phq9 sum-score 5-9)*<br>(12) |
| **PHQ15**<br>(14) | Patient Health Questionnaire: somatic symptoms module | Somatic Symptoms. | 2% high *(phq15 sum-score 15-30)*<br>17% medium *(phq15 sum-score 10-14)*<br>(14) |
| **PSS**<br>(15,16) | Perceived Stress Scale | Stress. | 1% high *(pss sum-score > 26)*<br>45% medium *(pss sum-score 14-26)*<br>(17,18) |
| **SIAS**<br>(19) | Social Interaction Anxiety Scale | Social Anxiety. | 16% *(sias sum-score >= 36)*<br>(20) |
| **STAI (Trait)**<br>(21) | State-Trait Anxiety Inventory P2 | Trait Anxiety. | 18% *(stai-trait sum-score >= 44)*<br>(22) |
| **WEMWBS**<br>(23) | Warwick-Edinburgh Mental Well-Being Scale | Mental Well-Being. | NA |
| **WHO5**<br>(24,25) | World Health Organisation-Five Well-Being Index | Well-Being. | 28% poor well-being *(who5 score: sum-score*4 <= 50)*<br>(25) |
| **WHOQOL**<br>(26) | World Health Organisation Quality of Life | Quality of Life: General, Social, Environmental, Physical, Psychological. | NA |

**Supplementary Table 2:** ***Cross-validated CCA signifying mental health functional implication of stomach-brain coupling.***

| | Stomach-Brain Coupling | | | Mental Health | | | Relationship | |
|---|---|---|---|---|---|---|---|---|
| | **Weight Stability** | **Explained Variance** | **nPCA** | **Weight Stability** | **Explained Variance** | **nPCA** | **In-Sample Correlation** | **Out-of-Sample Correlation** |
| **CCA Mode** | 0.221 | 0.714 | 71 | 0.328 | 33.733 | 1 | 0.886 | 0.323 |

The CCA mode characteristics of mental health stomach-brain coupling. This includes the 'weight stability' describing the stability of the CCA model via the average similarity (Pearson's correlation) of weights across each pair of training sets of the outer data splits. The percent 'explained variance' of the CCA model in contrast to the variance across the training sets of the outer data splits. The number of PCA components for each data type (nPCA) optimised for the significant CCA mode. The 'in-sample correlation' between the canonical variates of the training sets of the outer data splits. Finally, the 'out-of-sample correlation' of the canonical variates between the test sets of the outer data splits (cross-validated CCA relationship).

**Supplementary Table 3:** ***Relationship of Age and Gender with the Mental Health Associated Stomach-Brain Coupling CCA result.***

| | Age | Gender |
|---|---|---|
| CCA variate | Stomach-Brain Variate = r (197) = -0.040, p = 0.579<br>Mental-Health Variate = r (197) = -0.0129, p = 0.857 | Stomach-Brain Variate = t (197) = -0.546, p = 0.586<br>Mental Health Variate = t (197) = 0.664, p = 0.508 |

CCA with mental health scores and stomach-brain coupling was completed without the age and gender nuissance regressors. Age and gender effects were tested via a pearsons correlation with age and a t-test with gender with the subsequent CCA variate for each stomach-brain coupling and mental health.

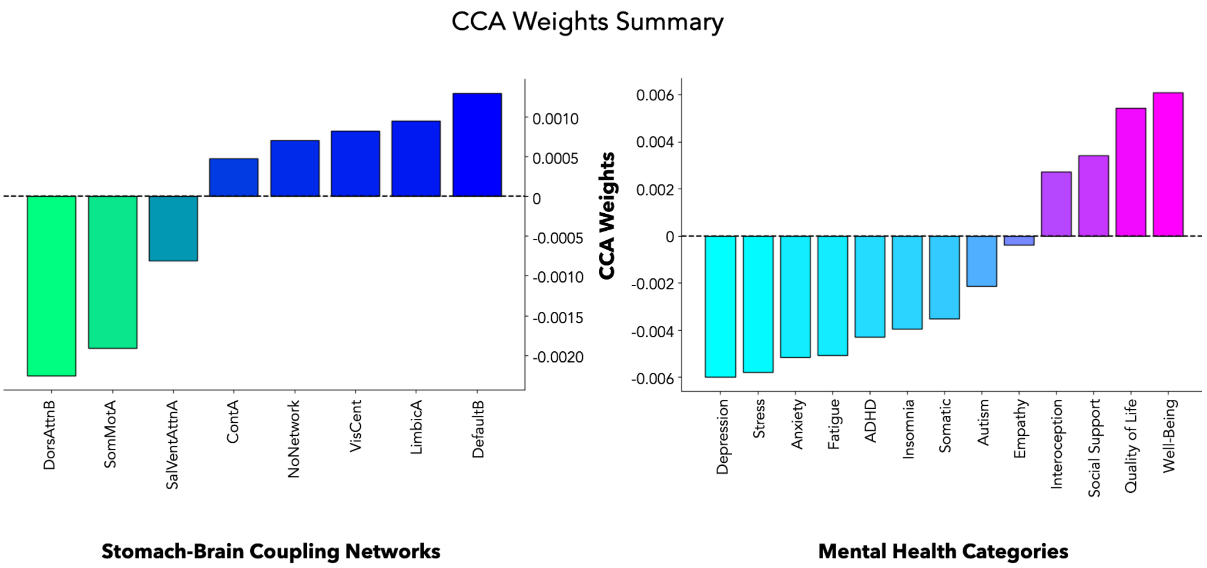


***Supplementary Figure 1*: CCA raw weights summary.**

Canonical weights from the mental health stomach-brain coupling CCA, summarised via averaging. Left shows the stomach-brain weights averaged according to yeo-7 networks (from left to right: DorsAttnB = Dorsal Attention B, SomMotA = Somatomotor A, SalVentAttnA = Salience Ventral Attention A, ContA = Control A, NoNetwork = no network assigned, VisCent = Visual A, LimbicA = Limbic A, DefaultB = Default Mode B) (48). Right illustrates the psychological weights averaged across mental health categories defined for visualisation in Figure 1.

**Supplementary Table 4:** ***Univariate correlation analysis of mental health PCA component and stomach-brain coupling for each DiFuMo parcellated region separately.*** Table shows significant regions of mental health stomach-brain coupling via univariate correlations which are largely consistent with the top stomach-brain coupling regions of the multivariate mental health CCA (see Supplementary Table 6).

| DiFuMo256 region label | Correlation with psych PCA component |
|---|---|
| Cingulate cortex mid-posterior | r = 0.164, p = 0.021 |
| Intraparietal sulcus anterior | r = 0.203, p = 0.004 |
| Superior frontal gyrus posterior LH | r = 0.173, p = 0.014 |
| Intraparietal sulcus posterior LH | r = 0.192, p = 0.007 |
| Angular gyrus superior LH | r = 0.207, p = 0.003 |
| Intermediate primus of Jensen RH | r = 0.198, p = 0.005 |
| Lateral fissure anterior | r = 0.152, p = 0.033 |
| Precentral sulcus inferior LH | r = 0.188, p = 0.008 |
| Superior occipital sulcus LH | r = 0.143, p = 0.044 |

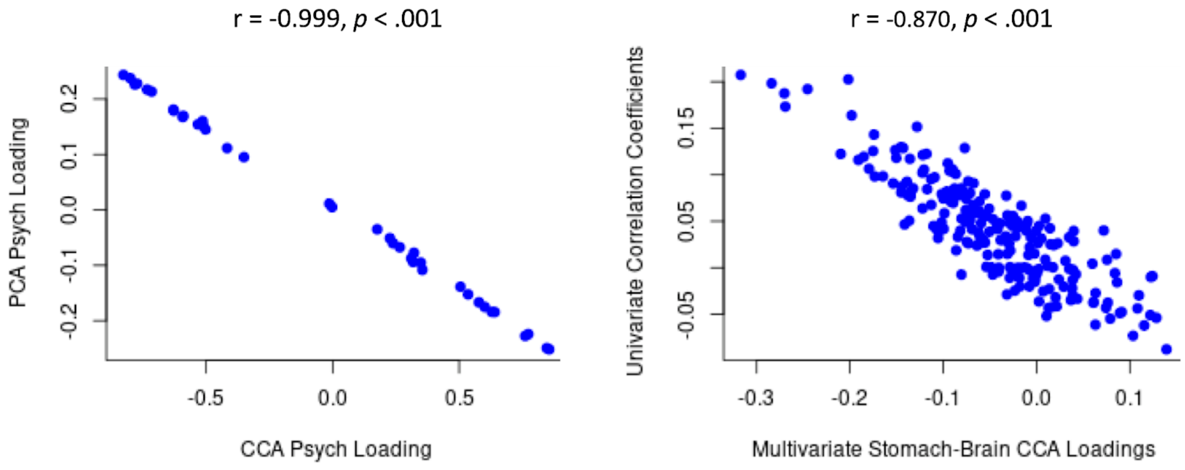


***Supplementary Figure 2*: Comparison of univariate correlation analysis with multivariate CCA analysis of stomach-brain coupling and mental health scores.**
Left is the highly correlated PCA component loadings of the mental health scores with the psychological loadings (structure correlations) from the CCA analysis, highlighting their similar structure. Right shows the individual univariate correlation coefficients of stomach-brain coupling in each of the 209 DiFuMo parcellated regions with the mental health PCA component, highly correlating with the stomach-brain CCA loadings of the multivariate CCA analysis.

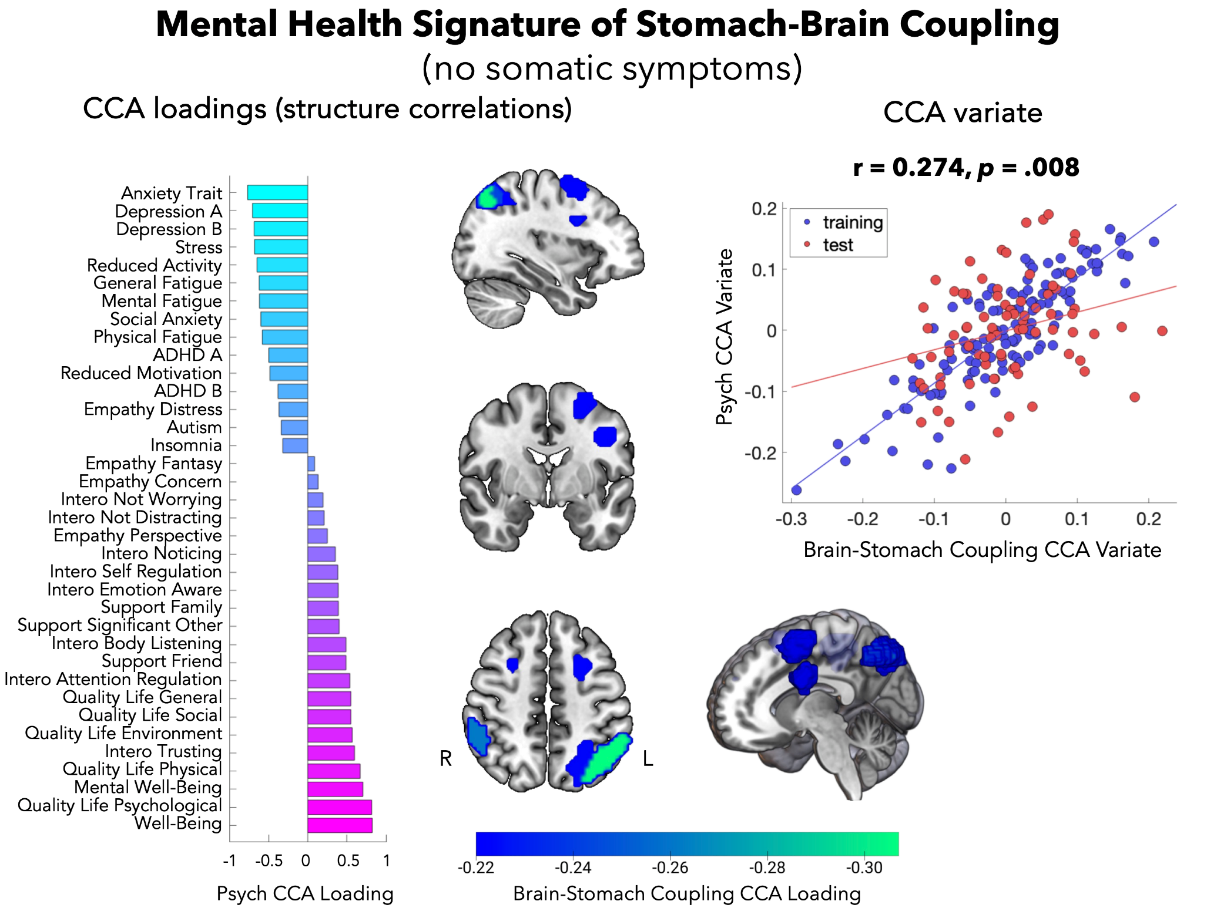


**Supplementary Figure 3: Mental health associated stomach-brain coupling result without somatic symptoms survey (PHQ15).**

Mental health signature of stomach-brain coupling CCA result persists and is highly similar without somatic symptoms survey (PHQ15), demonstrating that the mental health result is not driven by somatic or gastrointestinal symptoms.

**Supplementary Table 5:** ***EGG metric descriptive statistics in normogastric range.***

| Normogastric EGG Metric | Mean | Standard Deviation |
|---|---|---|
| Peak Frequency (Hz) | 0.050 | 0.004 |
| Maximum Power (μv2) | 245.104 | 562.098 |
| Proportion of Power | 0.394 | 0.204 |

**Supplementary Table 6:** ***Top 20 brain CCA loadings (structure correlations).***

| Brain region (DiFuMo256) | Network (Yeo7) | Canonical loading |
|---|---|---|
| Angular gyrus superior LH | ContA | -0.317 |
| Intermediate primus of Jensen RH | ContA | -0.284 |
| Precentral sulcus inferior LH | DorsAttnB | -0.270 |
| Superior frontal gyrus posterior LH | DorsAttnB | -0.269 |
| Intraparietal sulcus posterior LH | DorsAttnB | -0.245 |
| Posterior cingulate cortex middle | DefaultB | -0.210 |
| Intraparietal sulcus anterior | DorsAttnB | -0.202 |
| Cingulate cortex mid-posterior | SalVentAttnA | -0.198 |
| Middle frontal gyrus posterior RH | ContA | -0.191 |
| Superior parietal lobule posterior | DorsAttnB | -0.185 |
| Pericallosal sulcus middle | ContA | -0.179 |
| Superior parietal lobule middle | DorsAttnB | -0.175 |
| Superior occipital sulcus LH | DorsAttnB | -0.174 |
| Superior frontal gyrus posterior RH | DorsAttnB | -0.173 |
| Cingulate cortex mid-anterior | SalVentAttnA | -0.165 |
| Intraparietal sulcus LH | ContA | -0.154 |
| Middle temporal gyrus posterior | DorsAttnB | -0.151 |
| Intraparietal sulcus superior RH | DorsAttnB | -0.150 |
| Inferior frontal gyrus LH | ContA | -0.145 |
| Precentral sulcus superior | DorsAttnB | -0.145 |

ContA=frontoparietal control network, DorsAttnB=dorsal attention network, SalVentAttnA=salience ventral attention network.

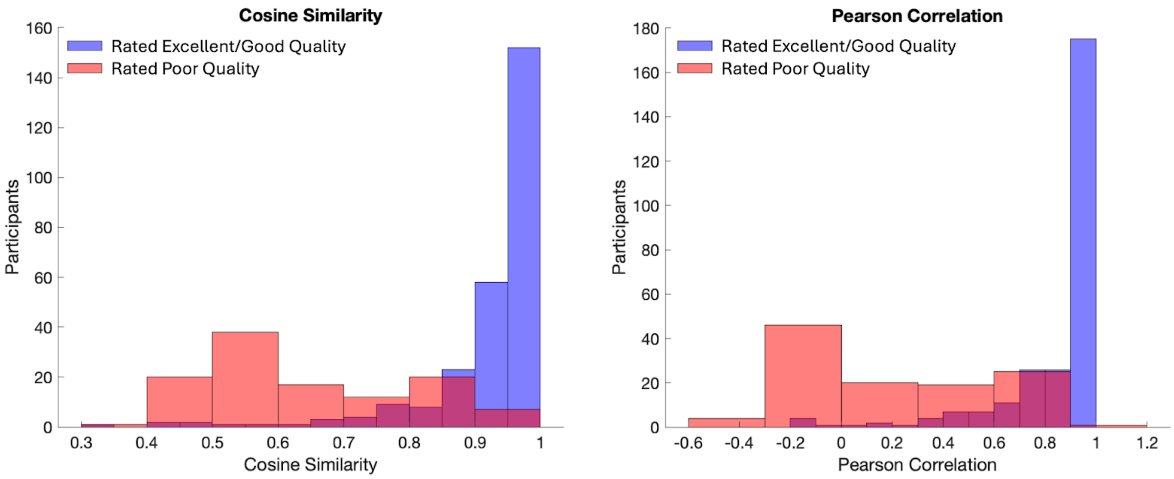


***Supplementary Figure 4*: Electrogastrography signal quality of included and excluded participants.**

EGG signal quality metrics (left: cosine similarity, right: Pearson correlation) of participants rated as ‘excellent/good quality’ in red and those rated as ‘poor quality’ in purple. These signal quality metrics were computed by comparing each individual's gastric FFT with the average FFT of 10 ideal participants with a very clear prominent normogastric peak.

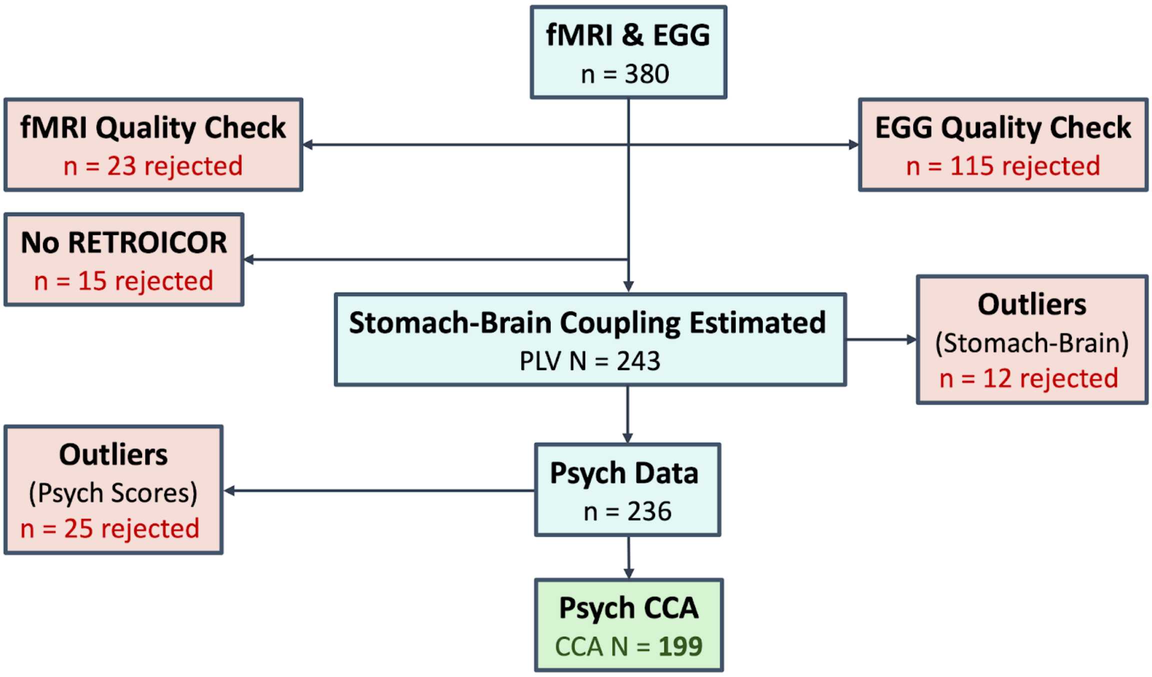


***Supplementary Figure 5*: Flow chart of analysis sample sizes from the Visceral Mind Project.**

Flow chart describing the sample size of each data type from the Visceral Mind Project, including resting-state fMRI, and electrogastrography (EGG) recordings. The neuroimaging and gastric physiological data was visually inspected resulting in low quality data rejections. Furthermore, data was rejected if there were corrupted cardiac or respiratory recordings which meant physiological noise could not be estimated via RETROICOR. The sample for estimating the stomach-brain coupling via phase-locking values (PLV) was 243. Outliers were rejected from both the stomach-brain coupling and mental health sampling scores. The final sample for the mental health functional correlate analysis with canonical correlation analysis (CCA) was 199.

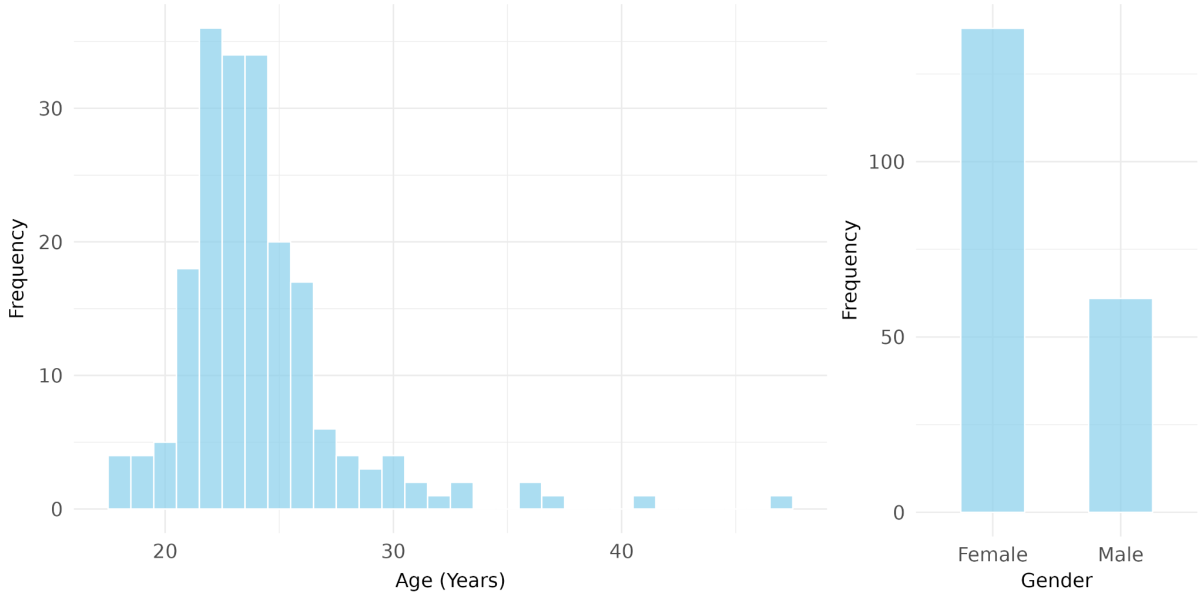


**Supplementary Figure 6: Demographic distributions for age and gender.**

Histograms of demographic data for the CCA sample: (left) frequency of age distribution, (right) frequency of gender.

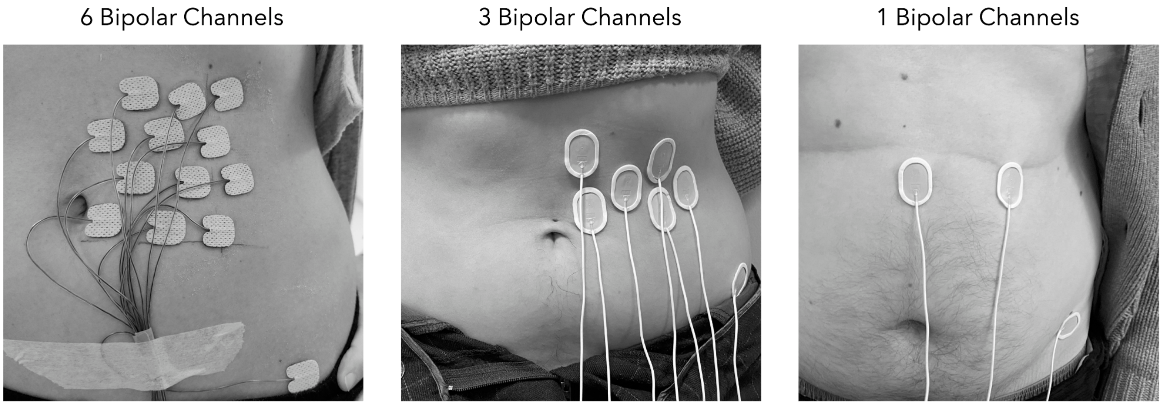


***Supplementary Figure 7*: Electrogastrography recording montages.**
Includes electrogastrography bipolar channel pairs (6, 3 or 1 recording pair) on left abdomen, with ground electrode on the left hip bone. For all montages, a single EGG channel pair was selected for stomach-brain coupling estimation.

**fMRIPrep preprocessing**

fMRI results included in this manuscript come from preprocessing performed using fMRIPrep 22.1.1 ((27,28); RRID:SCR_016216), which is based on Nipype 1.8.5 ((29); RRID:SCR_002502)

**fMRIPrep anatomical data preprocessing**

A total of 1 T1-weighted (T1w) images were found within the input BIDS dataset. The T1-weighted (T1w) image was corrected for intensity non-uniformity (INU) with N4BiasFieldCorrection (30), distributed with ANTs 2.3.3 ((31), RRID:SCR_004757), and used as T1w-reference throughout the workflow. The T1w-reference was then skull-stripped with a Nipype implementation of the antsBrainExtraction.sh workflow (from ANTs), using OASIS30ANTs as target template. Brain tissue segmentation of cerebrospinal fluid (CSF), white-matter (WM) and gray-matter (GM) was performed on the brain-extracted T1w using FAST (FSL 6.0.5.1:57b01774, RRID:SCR_002823, (32)). Brain surfaces were reconstructed using recon-all (FreeSurfer 7.2.0, RRID:SCR_001847, (33)), and the brain mask estimated previously was refined with a custom variation of the method to reconcile ANTs-derived and FreeSurfer-derived segmentations of the cortical gray-matter of Mindboggle (RRID:SCR_002438, (Klein et al., 2017)). Volume-based spatial normalization to two standard spaces (MNI152NLin2009cAsym, MNI152NLin6Asym) was performed through nonlinear registration with antsRegistration (ANTs 2.3.3), using brain-extracted versions of both T1w reference and the T1w template. The following templates were selected for spatial normalization: ICBM 152 Nonlinear Asymmetrical template version 2009c [(35), RRID:SCR_008796; TemplateFlow ID: MNI152NLin2009cAsym], FSL's MNI ICBM 152 non-linear 6th Generation Asymmetric Average Brain Stereotaxic Registration Model [(36), RRID:SCR_002823; TemplateFlow ID: MNI152NLin6Asym].

**fMRIPrep functional data preprocessing**

For each of the 3 BOLD runs found per subject (across all tasks and sessions), the following preprocessing was performed. First, a reference volume and its skull-stripped version were generated using a custom methodology of fMRIPrep. Head-motion parameters with respect to the BOLD reference (transformation matrices, and six corresponding rotation and translation parameters) are estimated before any spatiotemporal filtering using mcflirt(FSL 6.0.5.1:57b01774, (37)). BOLD runs were slice-time corrected to 0.658s (0.5 of slice acquisition range 0s-1.31s) using 3dTshift from AFNI ((38), RRID:SCR_005927). The BOLD time-series (including slice-timing correction when applied) were resampled onto their original, native space by applying the transforms to correct for head-motion. These resampled BOLD time-series will be referred to as preprocessed BOLD in original space, or just preprocessed BOLD. The BOLD reference was then co-registered to the T1w reference using bbregister (FreeSurfer) which implements boundary-based registration (39). Co-registration was configured with six degrees of freedom. Several confounding time-series were calculated based on the preprocessed BOLD: framewise displacement (FD), DVARS and three region-wise global signals. FD was computed using two formulations following Power (absolute sum of relative motions, (40)) and Jenkinson (relative root mean square displacement between affines, (37)). FD and DVARS are calculated for each functional run, both using their implementations in Nipype (following the definitions by (40)). The three global signals are extracted within the CSF, the WM, and the whole-brain masks. Additionally, a set of physiological regressors were extracted to allow for component-based noise correction (CompCor, (Behzadi et al., 2007)). Principal components are estimated after high-pass filtering the preprocessed BOLD time-series (using a discrete cosine filter with 128s cut-off) for the two CompCor variants: temporal (tCompCor) and anatomical (aCompCor). tCompCor components are then calculated from the top 2% variable voxels within the brain mask. For aCompCor, three probabilistic masks (CSF, WM and combined CSF+WM) are generated in anatomical space. The implementation differs from that of Behzadi et al. in that instead of eroding the masks by 2 pixels on BOLD space, a mask of pixels that likely contain a volume fraction of GM is subtracted from the aCompCor masks. This mask is obtained by dilating a GM mask extracted from the FreeSurfer's aseg segmentation, and it ensures components are not extracted from voxels containing a minimal fraction of GM. Finally, these masks are resampled into BOLD space and binarized by thresholding at 0.99 (as in the original implementation). Components are also calculated separately within the WM and CSF masks. For each CompCor decomposition, the k components with the largest singular values are retained, such that the retained components' time series are sufficient to explain 50 percent of variance across the nuisance mask (CSF, WM, combined, or temporal). The remaining components are dropped from consideration. The head-motion estimates calculated in the correction step were also placed within the corresponding confounds file. The confound time series derived from head motion estimates and global signals were expanded with the inclusion of temporal derivatives and quadratic terms for each (42). Frames that exceeded a threshold of 0.5 mm FD or 1.5 standardized DVARS were annotated as motion outliers. Additional nuisance timeseries are calculated by means of principal components analysis of the signal found within a thin band (crown) of voxels around the edge of the brain, as proposed by (43). The BOLD time-series were resampled into standard space, generating a preprocessed BOLD run in MNI152NLin2009cAsym space. First, a reference volume and its skull-stripped version were generated using a custom methodology of fMRIPrep. The BOLD time-series were resampled onto the following surfaces (FreeSurfer reconstruction nomenclature): fsaverage. Grayordinates files (44) containing 91k samples were also generated using the highest-resolution fsaverage as intermediate standardized surface space. All resamplings can be performed with a single interpolation step by composing all the pertinent transformations (head-motion transform matrices, susceptibility distortion correction when available, and co-registrations to anatomical and output spaces). Gridded (volumetric) resamplings were performed using antsApplyTransforms (ANTs), configured with Lanczos interpolation to minimize the smoothing effects of other kernels (45). Non-gridded (surface) resamplings were performed using mri_vol2surf (FreeSurfer).

Many internal operations of fMRIPrep use Nilearn 0.9.1 ((46), RRID:SCR_001362), mostly within the functional processing workflow. For more details of the pipeline, see (47).

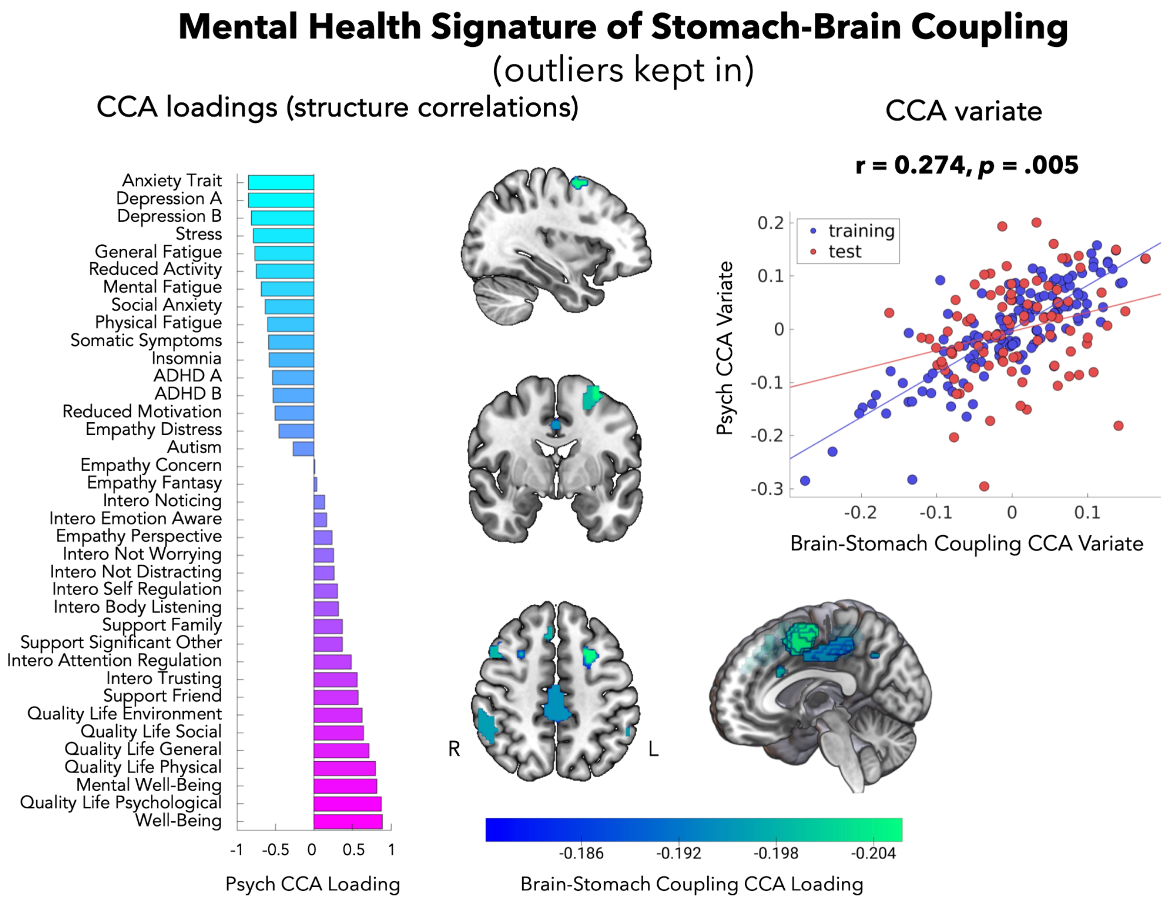


**Supplementary Figure 8: Mental Health Associated Stomach-Brain Coupling CCA result with psychiatric outliers kept in.**

CCA of multidimensional mental health and stomach-brain coupling with psychiatric outliers kept in, as requested by a reviewer.

### References


1. Allison C, Auyeung B, Baron-Cohen S. Toward Brief "Red Flags" for Autism Screening: The Short Autism Spectrum Quotient and the Short Quantitative Checklist in 1,000 Cases and 3,000 Controls. J Am Acad Child Adolesc Psychiatry. 2012 Feb 1;51(2):202-212.e7.
2. Baron-Cohen S, Wheelwright S, Skinner R, Martin J, Clubley E. The Autism-Spectrum Quotient (AQ): Evidence from Asperger Syndrome/High-Functioning Autism, Males and Females, Scientists and Mathematicians. J Autism Dev Disord. 2001 Feb 1;31(1):5–17.
3. Kessler RC, Adler L, Ames M, Demler O, Faraone S, Hiripi E, et al. The World Health Organization adult ADHD self-report scale (ASRS): a short screening scale for use in the general population. Psychol Med. 2005 Feb;35(2):245–56.
4. Davis MH. Interpersonal Reactivity Index [Internet]. 1980 [cited 2023 Nov 17]. Available from: http://doi.apa.org/getdoi.cfm?doi=10.1037/t01093-000
5. Davis MH. Measuring individual differences in empathy: Evidence for a multidimensional approach. J Pers Soc Psychol. 1983;44(1):113–26.
6. Morin CM, Belleville G, Bélanger L, Ivers H. The Insomnia Severity Index: Psychometric Indicators to Detect Insomnia Cases and Evaluate Treatment Response. Sleep. 2011 May 1;34(5):601–8.
7. Mehling WE, Price C, Daubenmier JJ, Acree M, Bartmess E, Stewart A. The Multidimensional Assessment of Interoceptive Awareness (MAIA). PLOS ONE. 2012 Nov 1;7(11):e48230.
8. Bech P, Timmerby N, Martiny K, Lunde M, Soendergaard S. Psychometric evaluation of the Major Depression Inventory (MDI) as depression severity scale using the LEAD (Longitudinal Expert Assessment of All Data) as index of validity. BMC Psychiatry. 2015 Aug 5;15(1):190.
9. Olsen LR, Jensen DV, Noerholm V, Martiny K, Bech P. The internal and external validity of the Major Depression Inventory in measuring severity of depressive states. Psychol Med. 2003 Feb;33(2):351–6.
10. Smets EMA, Garssen B, Bonke B, De Haes JCJM. The multidimensional Fatigue Inventory (MFI) psychometric qualities of an instrument to assess fatigue. J Psychosom Res. 1995 Apr 1;39(3):315–25.
11. Zimet GD, Dahlem NW, Zimet SG, Farley GK. The Multidimensional Scale of Perceived Social Support. J Pers Assess. 1988 Mar 1;52(1):30–41.
12. Kroenke K, Spitzer RL, Williams JBW. The PHQ-9. J Gen Intern Med. 2001;16(9):606–13.
13. Spitzer RL, Kroenke K, Williams JBW, and the Patient Health Questionnaire Primary Care Study Group. Validation and Utility of a Self-report Version of PRIME-MDThe PHQ Primary Care Study. JAMA. 1999 Nov 10;282(18):1737–44.
14. Kroenke K, Spitzer RL, Williams JBW. The PHQ-15: Validity of a New Measure for Evaluating the Severity of Somatic Symptoms. Psychosom Med. 2002 Apr;64(2):258.
15. Cohen S. Perceived stress in a probability sample of the United States. In: The social psychology of health. Thousand Oaks, CA, US: Sage Publications, Inc; 1988. p. 31–67. (The Claremont Symposium on Applied Social Psychology).

16. Cohen S, Kamarck T, Mermelstein R. A Global Measure of Perceived Stress. J Health Soc Behav. 1983;24(4):385–96.
17. Philpott LF, Leahy-Warren P, FitzGerald S, Savage E. Prevalence and associated factors of paternal stress, anxiety, and depression symptoms in the early postnatal period. Glob Ment Health. 2022 Jan;9:306–21.
18. Swaminathan A, Viswanathan S, Gnanadurai T, Ayyavoo S, Manickam T. Perceived stress and sources of stress among first-year medical undergraduate students in a private medical college - Tamil Nadu. Natl J Physiol Pharm Pharmacol. 2016;6(1):9–14.
19. Mattick RP, Clarke JC. Development and validation of measures of social phobia scrutiny fear and social interaction anxiety11Editor's note: This article was written before the development of some contemporary measures of social phobia, such as the Social Phobia and Anxiety Inventory (Turner et al., 1989). We have invited this article for publication because of the growing interest in the scales described therein. S.T. Behav Res Ther. 1998 Apr 1;36(4):455–70.
20. Peters L. Discriminant validity of the Social Phobia and Anxiety Inventory (SPAI), the Social Phobia Scale (SPS) and the Social Interaction Anxiety Scale (SIAS). Behav Res Ther. 2000 Sep 1;38(9):943–50.
21. Spielberger CD. State-Trait Anxiety Inventory for Adults [Internet]. APA PsycTests.; 1983 [cited 2023 Nov 22]. Available from: http://doi.apa.org/getdoi.cfm?doi=10.1037/t06496-000
22. Ercan İ, Hafizoglu S, Özkaya G, Kirli S, Yalcintas E, Akkaya C. EXAMINING CUT-OFF VALUES FOR THE STATE-TRAIT ANXIETY INVENTORY. Rev Argent Clin Psicol [Internet]. 2015 [cited 2023 Dec 7];24(2). Available from: https://avesis.uludag.edu.tr/yayin/9e6b417b-a269-4659-a47b-21ca9d7a352e/examining-cut-off-values-for-the-state-trait-anxiety-inventory
23. Tennant R, Hiller L, Fishwick R, Platt S, Joseph S, Weich S, et al. The Warwick-Edinburgh Mental Well-being Scale (WEMWBS): development and UK validation. Health Qual Life Outcomes. 2007 Nov 27;5(1):63.
24. Bech P, Olsen LR, Kjoller M, Rasmussen NK. Measuring well-being rather than the absence of distress symptoms: a comparison of the SF-36 Mental Health subscale and the WHO-Five well-being scale. Int J Methods Psychiatr Res. 2003;12(2):85–91.
25. Topp CW, Østergaard SD, Søndergaard S, Bech P. The WHO-5 Well-Being Index: A Systematic Review of the Literature. Psychother Psychosom. 2015 Mar 28;84(3):167–76.
26. Skevington SM, Lotfy M, O'Connell KA. The World Health Organization's WHOQOL-BREF quality of life assessment: Psychometric properties and results of the international field trial. A Report from the WHOQOL Group. Qual Life Res. 2004 Mar 1;13(2):299–310.
27. Esteban O, Markiewicz CJ, Blair RW, Moodie CA, Isik AI, Erramuzpe A, et al. fMRIPrep: a robust preprocessing pipeline for functional MRI. Nat Methods. 2019 Jan;16(1):111–6.
28. Esteban O, Markiewicz CJ, Goncalves M, Provins C, Kent JD, DuPre E, et al. fMRIPrep: a robust preprocessing pipeline for functional MRI [Internet]. Zenodo; 2022 [cited 2023 Feb 14]. Available from: https://zenodo.org/record/7430291
29. Gorgolewski K, Ziegler E, Ellis DG, Notter MP, Perkins LN. Nipype. 2018.

30. Tustison NJ, Avants BB, Cook PA, Zheng Y, Egan A, Yushkevich PA, et al. N4ITK: Improved N3 Bias Correction. IEEE Trans Med Imaging. 2010 Jun;29(6):1310–20.
31. Avants BB, Epstein CL, Grossman M, Gee JC. Symmetric diffeomorphic image registration with cross-correlation: Evaluating automated labeling of elderly and neurodegenerative brain. Med Image Anal. 2008 Feb 1;12(1):26–41.
32. Zhang Y, Brady M, Smith S. Segmentation of brain MR images through a hidden Markov random field model and the expectation-maximization algorithm. IEEE Trans Med Imaging. 2001 Jan;20(1):45–57.
33. Dale AM, Fischl B, Sereno MI. Cortical Surface-Based Analysis: I. Segmentation and Surface Reconstruction. NeuroImage. 1999 Feb 1;9(2):179–94.
34. Klein A, Ghosh SS, Bao FS, Giard J, Häme Y, Stavsky E, et al. Mindboggling morphometry of human brains. PLOS Comput Biol. 2017 Feb 23;13(2):e1005350.
35. Fonov V, Evans A, McKinstry R, Almli C, Collins D. Unbiased nonlinear average age-appropriate brain templates from birth to adulthood. NeuroImage. 2009 Jul 1;47:S102.
36. Evans AC, Janke AL, Collins DL, Baillet S. Brain templates and atlases. NeuroImage. 2012 Aug 15;62(2):911–22.
37. Jenkinson M, Bannister P, Brady M, Smith S. Improved Optimization for the Robust and Accurate Linear Registration and Motion Correction of Brain Images. NeuroImage. 2002 Oct 1;17(2):825–41.
38. Cox RW, Hyde JS. Software tools for analysis and visualization of fMRI data. NMR Biomed. 1997;10(4–5):171–8.
39. Greve DN, Fischl B. Accurate and robust brain image alignment using boundary-based registration. NeuroImage. 2009 Oct 15;48(1):63–72.
40. Power JD, Mitra A, Laumann TO, Snyder AZ, Schlaggar BL, Petersen SE. Methods to detect, characterize, and remove motion artifact in resting state fMRI. NeuroImage. 2014 Jan 1;84:320–41.
41. Behzadi Y, Restom K, Liau J, Liu TT. A component based noise correction method (CompCor) for BOLD and perfusion based fMRI. NeuroImage. 2007 Aug 1;37(1):90–101.
42. Satterthwaite TD, Elliott MA, Gerraty RT, Ruparel K, Loughead J, Calkins ME, et al. An improved framework for confound regression and filtering for control of motion artifact in the preprocessing of resting-state functional connectivity data. NeuroImage. 2013 Jan 1;64:240–56.
43. Patriat R, Reynolds RC, Birn RM. An improved model of motion-related signal changes in fMRI. NeuroImage. 2017 Jan 1;144:74–82.
44. Glasser MF, Sotiropoulos SN, Wilson JA, Coalson TS, Fischl B, Andersson JL, et al. The minimal preprocessing pipelines for the Human Connectome Project. NeuroImage. 2013 Oct 15;80:105–24.
45. Lanczos C. Evaluation of Noisy Data. J Soc Ind Appl Math Ser B Numer Anal. 1964 Jan;1(1):76–85.
46. Abraham A, Pedregosa F, Eickenberg M, Gervais P, Mueller A, Kossaifi J, et al. Machine learning for neuroimaging with scikit-learn. Front Neuroinformatics [Internet]. 2014 [cited 2023 Feb 14];8. Available from: https://www.frontiersin.org/articles/10.3389/fninf.2014.00014
47. Preprocessing pipeline details - fmriprep version documentation [Internet].

Available from: https://fmriprep.readthedocs.io/en/latest/workflows.html

48. Yeo BTT, Krienen FM, Sepulcre J, Sabuncu MR, Lashkari D, Hollinshead M, et al. The organization of the human cerebral cortex estimated by intrinsic functional connectivity. J Neurophysiol. 2011;106(3):1125–65.
